# CD59 organizes the plasma membrane to sustain oncogenic Ras-MAPK signaling and is a targetable vulnerability in acute myeloid leukemia

**DOI:** 10.64898/2026.08.11.744285

**Authors:** Abdula Z. Maher, Dhanoop Manikoth Ayyathan, Severine Cathelin, Paige Roehrig, Stephanie Z. Liu, Yitong Yang, Alex C.H. Liu, Mohsen Hosseini, Elise Quadri, Tiffany Villeneuve, Simran Kaur, Vivian Wang, Erwin M. Schoof, Andrea Arruda, Mark D. Minden, Christopher B. Marshall, Aaron Schimmer, Stephanie Xie, John E. Dick, Steven M. Chan

## Abstract

Acute myeloid leukemia (AML) is a clinically heterogeneous disease. Although the genetic abnormalities associated with poor prognosis are well defined, how they drive unfavorable outcomes remains unclear. Using published gene-expression and dependency datasets, we searched for cell-surface protein-coding genes associated with poor survival and required for AML growth, prioritizing this class of proteins for its accessibility to biologics. This search identified CD59, a GPI-anchored protein with a canonical role in complement regulation, whose high mRNA expression correlates with adverse-risk genetics and stemness signatures. *CD59* silencing impaired proliferation across genetically diverse AML cell lines, reduced leukemic burden, and extended survival in cell xenograft models. Moreover, CD59 expression was enriched on leukemic stem cells (LSCs), and its depletion impaired LSC self-renewal and primary AML engraftment *in vivo* while sparing normal hematopoiesis. Mechanistically, these effects reflected a non-canonical role for CD59 in sustaining Ras-MAPK signaling, whereby its loss depleted inner-leaflet phosphatidylserine and impaired Ras and c-Raf membrane recruitment and activation. rILYd4, a recombinant fragment of the bacterial toxin intermedilysin that binds and degrades CD59, recapitulated these effects and sensitized cells to venetoclax in vivo. These findings reveal CD59 as a critical regulator of Ras-MAPK signaling required for AML growth and nominate its rILYd4-mediated degradation as a therapeutic strategy.

## Introduction

Acute myeloid leukemia (AML) is a hematologic malignancy arising from the clonal expansion of myeloid progenitors with impaired differentiation^1,2^. Its clinical course is highly heterogeneous. Although some patients achieve durable remissions, those with adverse-risk AML, defined by the presence of specific genetic abnormalities including complex cytogenetics, *TP53* mutations, and myelodysplasia-related gene mutations, have a median survival of less than one year despite aggressive treatments with intensive chemotherapy and allogeneic stem cell transplantation^3–5^. These cytogenetic and molecular abnormalities are well-established prognostic markers, but the mechanisms by which they drive poor outcomes remain incompletely understood^6^. Defining the therapeutic targets that mediate these mechanisms can lead to treatments that improve survival in adverse-risk AML.

Among potential therapeutic targets, cell-surface proteins are especially attractive because their extracellular domains are directly accessible to biologics. Antibodies, antibody-drug conjugates, and engineered protein binders can achieve specificity that small molecules rarely attain against intracellular targets, thereby minimizing off-target effects. Accessibility from outside the cell also removes the requirement for membrane permeability and extends the druggable proteome to adhesion molecules, transporters, and glycosylphosphatidylinositol (GPI)-anchored proteins that frequently lack the defined pockets required for small-molecule binding. Because biologics engage extended surfaces rather than such pockets, these proteins remain tractable even where small molecules fail.

In AML, this potential remains largely unrealized. Gemtuzumab ozogamicin, a CD33-directed antibody-drug conjugate, is the only surface antigen-directed agent approved for the disease^7^. Its benefit is restricted to favorable and intermediate cytogenetic risk groups and is not evident in adverse-risk AML^8^. Its use is further limited by hepatotoxicity and prolonged myelosuppression^9,10^. Myelosuppression in particular reflects a common drawback of the mechanism by which most surface proteins are currently targeted. In the case of gemtuzumab, CD33 is used to direct a cytotoxic payload (calicheamicin), so cells are killed based on antigen expression^11^. Because CD33 is also expressed on normal myeloid progenitor and differentiated cells, the therapeutic window depends on a narrow difference in antigen expression^12^. The search for targets with leukemia-restricted expression has yielded few candidates, as most myeloid surface antigens are shared with normal hematopoiesis. An alternative approach is to select targets by dependency rather than expression and to suppress their function rather than deliver cytotoxicity. A surface protein required by AML cells but dispensable in normal tissues can be targeted even when both compartments express it, because only the dependent cells are affected by its loss.

Based on the above considerations, we searched for a surface protein target that fulfills two criteria: 1) its expression in AML is associated with poor clinical outcomes, identifying a candidate that may functionally drive an aggressive phenotype; and 2) it is essential for AML but less essential for normal tissues, providing a therapeutic window based on dependency rather than differential expression. This search led us to CD59, a GPI-anchored cell-surface protein whose canonical function is to regulate the complement system by blocking assembly of the membrane attack complex (MAC)^13^. In this study, we show that it sustains

AML proliferation and survival through a mechanism independent of its role in complement regulation, acting instead through modulation of plasma membrane composition and intracellular signaling. To translate this into a therapeutic strategy, we leveraged a naturally occurring bacterial protein fragment that binds CD59 and routes it to the lysosome, clearing it from the cell surface and suppressing AML growth. Together, our findings establish CD59 as a functionally required surface vulnerability whose clearance reduces AML growth and demonstrate a targeted degradation strategy with therapeutic potential.

## Results

### Increased *CD59* expression is associated with poor outcomes, adverse-risk genetics, and stemness in AML

To identify surface proteins associated with outcome in AML, we analysed patients from the BeatAML2 cohort with a diagnosis of AML, diagnostic RNA-seq, and survival data (n=440)^14^. For each of 371 cluster of differentiation (CD) genes encoding surface proteins, we fitted a univariable Cox proportional hazards model to estimate the hazard ratio (HR) for overall survival per standard deviation increase in z-score normalized expression, with p-values adjusted using the Benjamini-Hochberg procedure. This analysis identified 14 genes with an HR > 1.2 and adjusted p-value < 0.05 (**Fig. 1A**). As an alternative analytical approach, we compared the expression of CD genes between patients with poor outcomes (defined as survival < 1 year) (n=78) versus favorable outcomes (survival > 3 years) (n=183). This approach identified three genes with a significant increase in expression in the poor-outcome cohort (ΔZ-score > 0.5 and adjusted P-value < 0.05) (**Fig. 1A**). The three genes, *CD59*, *CD109*, and *MME*, were also identified using the first analytical approach (**Fig. 1A**).

**Figure 1:**
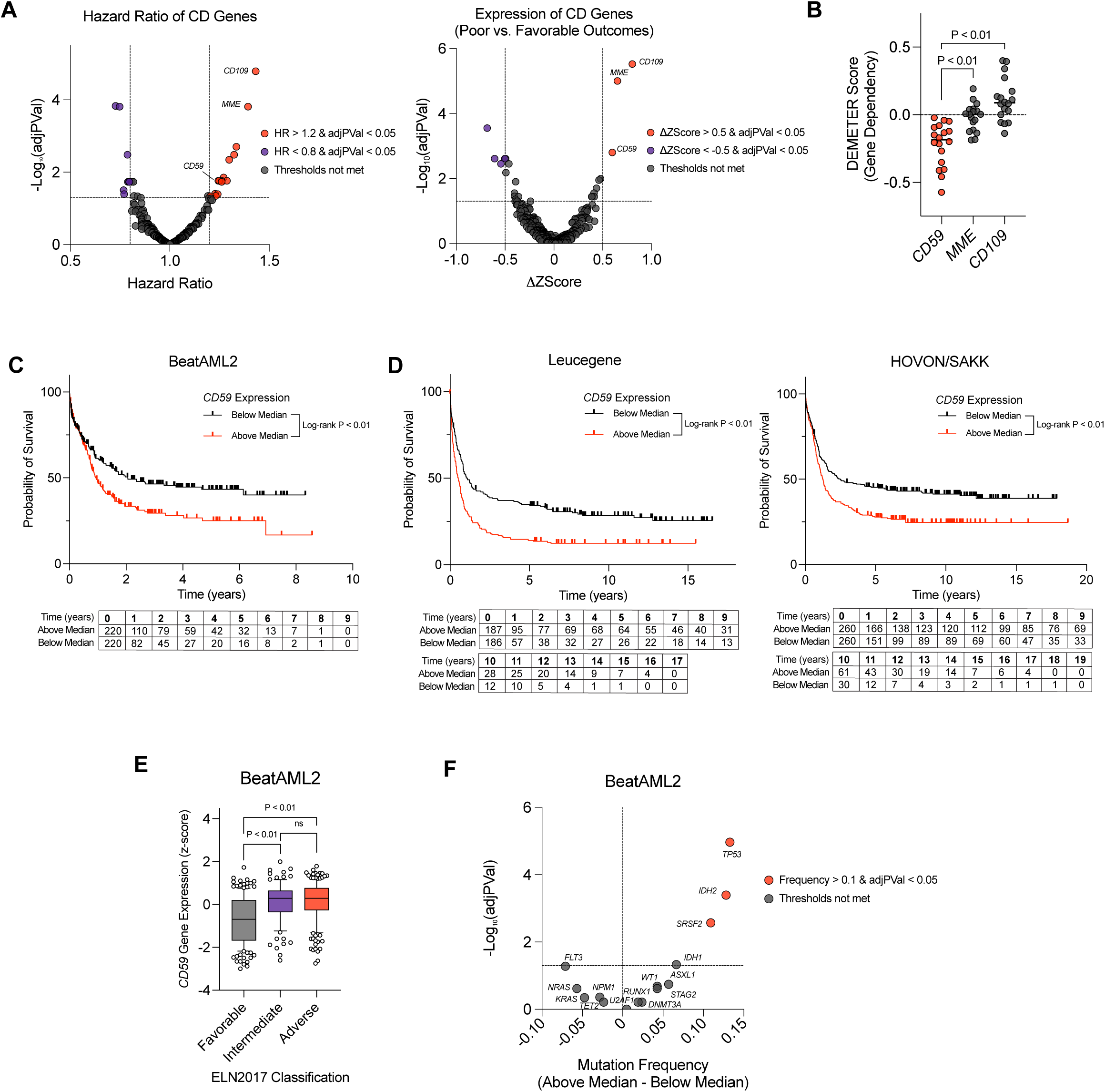
*CD59* is a prognostic marker and functionally required cell-surface antigen in AML. **(A)** Identification of cluster-of-differentiation (CD) genes associated with overall survival in the BeatAML2 cohort (n = 440 patients with AML, diagnostic RNA-seq, and survival data). Left: volcano plot of univariable Cox proportional-hazards results for 371 CD genes, showing the hazard ratio for overall survival per standard-deviation increase in z-scored expression (x-axis) versus −log₁₀ Benjamini-Hochberg-adjusted P value (y-axis). Red, HR > 1.2 and adjusted P < 0.05; blue, HR < 0.8 and adjusted P < 0.05; grey, thresholds not met. Right: differential expression of the 371 CD genes between patients with poor outcome (overall survival < 1 year, n = 78) and favorable outcome (overall survival > 3 years, n = 183), showing the difference in mean z-score (ΔZ-score; poor − favorable) versus −log₁₀ adjusted P value. Red, ΔZ-score > 0.5 and adjusted P < 0.05; blue, ΔZ-score < −0.5 and adjusted P < 0.05. **(B)** Gene-dependency (DEMETER2 RNAi) scores for *CD59*, *MME*, and *CD109* across 18 AML cell lines in DepMap. **(C)** Kaplan-Meier overall-survival curves for patients in BeatAML2 (n = 440) stratified by *CD59* expression above (red) versus below (black) the cohort median. P value, log-rank test. **(D)** Kaplan-Meier overall-survival analysis as in (C) for two independent validation cohorts: Leucegene (n = 373) and HOVON/SAKK (n = 520). *CD59* above (red) versus below (black) median; P values, log-rank test; numbers at risk below each curve. **(E)** *CD59* expression (z-score) in BeatAML2 patients grouped by 2017 European LeukemiaNet (ELN2017) risk classification: Favorable, Intermediate, and Adverse. Boxes show median and interquartile range; points are individual patients. **(F)** Enrichment of recurrent AML driver mutations in *CD59*-above-median versus *CD59*-below-median patients (BeatAML2). Volcano plot of the difference in mutation frequency (above − below median; x-axis) versus −log₁₀ adjusted P value (y-axis); Fisher’s exact test with Benjamini-Hochberg correction. Red, frequency difference > 0.1 and adjusted P < 0.05; grey, thresholds not met.

To determine which of these genes are functionally required for AML, we examined their dependency scores across 18 AML cell lines in DepMap. This analysis showed that *CD59* exhibited the greatest dependency of the three genes (**Fig. 1B**). We next asked whether loss of *CD59* is likely tolerated by normal cells. *CD59* is not classified as a common essential gene in DepMap; individuals and mouse models with biallelic CD59 loss-of-function mutations are viable with preserved hematopoiesis; and *PIGA*-mutant stem cell clones lacking CD59 sustain long-term multilineage blood production in paroxysmal nocturnal hemoglobinuria^15–18^. CD59 therefore appeared to be required by AML cells yet dispensable in normal hematopoiesis. This differential dependency, together with its surface accessibility and the correlation between its high expression and poor clinical outcomes, led us to prioritize CD59 for further evaluation.

We first validated the clinical relevance of *CD59* in AML using independent datasets. In the BeatAML2 cohort, patients with *CD59* expression above the median (*CD59*^HIGH^) had significantly worse overall survival than those below the median (*CD59*^LOW^) (median OS = 11.24 months, 95% CI [9.2 months to 14.42 months] versus 24.67 months, 95% CI [17.08 months to 73.63 months]) by Kaplan-Meier analysis (**Fig. 1C**). These associations were reproduced in two independent AML cohorts, Leucegene (n=373) and HOVON/SAKK (n=520)^19,20^ (**Fig. 1D**). We next asked whether *CD59* expression correlated with established markers of treatment response and prognostic risk. *CD59* expression was higher in patients with primary refractory disease than in those achieving a remission (**Extended Fig. 1A**) and in patients with adverse- and intermediate-risk disease by ELN2017 criteria compared with those with favourable-risk disease (**Fig. 1E**). In addition, *CD59*^HIGH^ patients were enriched for mutations that confer poor prognosis, notably *TP53* and *SRSF2* (**Fig. 1F**). Interestingly, in a multivariable model adjusting for ELN2017 risk, age, and AML ontogeny (de novo versus secondary), *CD59* expression did not retain independent prognostic significance (**Extended Fig. 1B**). This lack of independence suggests that CD59 does not act separately from these poor-risk features but is tightly coupled to them, possibly mediating the aggressive biology they confer.

A common biological feature underlying poor prognosis in AML is the persistence of cells with stem cell properties, referred to as leukemic stem cells (LSCs)^21,22^. We therefore asked whether AML with high *CD59* expression is enriched for stem cell transcriptional programs. The LSC17 score is a signature derived from functionally defined LSC fractions that predicts adverse outcomes in AML^23^. Across four independent cohorts (BeatAML2, Leucegene, HOVON/SAKK, and TCGA-AML), *CD59*^HIGH^ patients had higher LSC17 scores than *CD59*^LOW^ patients (**Extended Fig. 1C**). To determine whether *CD59* expression tracks with the differentiation state of the leukemic hierarchy, we scored each sample in the BeatAML2 cohort for seven transcriptional signatures spanning leukemic stem and progenitor cells (LSPCs) to more mature leukemic populations^24^. *CD59*^HIGH^ samples showed higher scores for immature states (LSPC-Quiescent, LSPC-Primed, LSPC-Cycle, and GMP-like), whereas *CD59*^LOW^ samples showed higher scores for the more differentiated ProMono-like, Mono-like, and cDC-like states (**Extended Fig. 1D**). Together, these data indicate that high *CD59* expression is associated with poor outcomes, adverse-risk prognostic markers, and a primitive, stemness-high leukemic state, prompting us to examine its functional role in AML.

### Silencing *CD59* impairs AML proliferation and viability independent of complement activity

To investigate the functional role of CD59 in AML, we tested the impact of *CD59* knockdown across a genetically diverse panel of nine AML cell lines spanning a range of baseline *CD59* mRNA and protein expression (**Extended Fig. 2A; Extended Table 1**). Each line was transduced with lentiviral vectors encoding one of two independent short hairpin RNAs targeting *CD59* (sh*CD59*-1, sh*CD59*-2) or a non-targeting control (shNT), expressed from a doxycycline-inducible promoter and linked to a blue fluorescent protein (BFP) reporter. Transduced (BFP⁺) cells were mixed 1:1 with untransduced (BFP⁻) cells, and knockdown was induced with 100 ng/ml doxycycline (**Extended Fig. 2B**). Longitudinal tracking of the BFP⁺ fraction over 10 days showed that sh*CD59*-expressing cells were progressively depleted in all nine lines, whereas shNT-expressing cells maintained relatively stable representation (**Fig. 2A**; **Extended Fig. 2C**). Both hairpins produced concordant phenotypes in every line. Moreover, baseline *CD59* mRNA expression correlated positively with the magnitude of growth impairment following knockdown (**Extended Fig. 2D**). These findings are consistent with an on-target effect.

**Figure 2.**
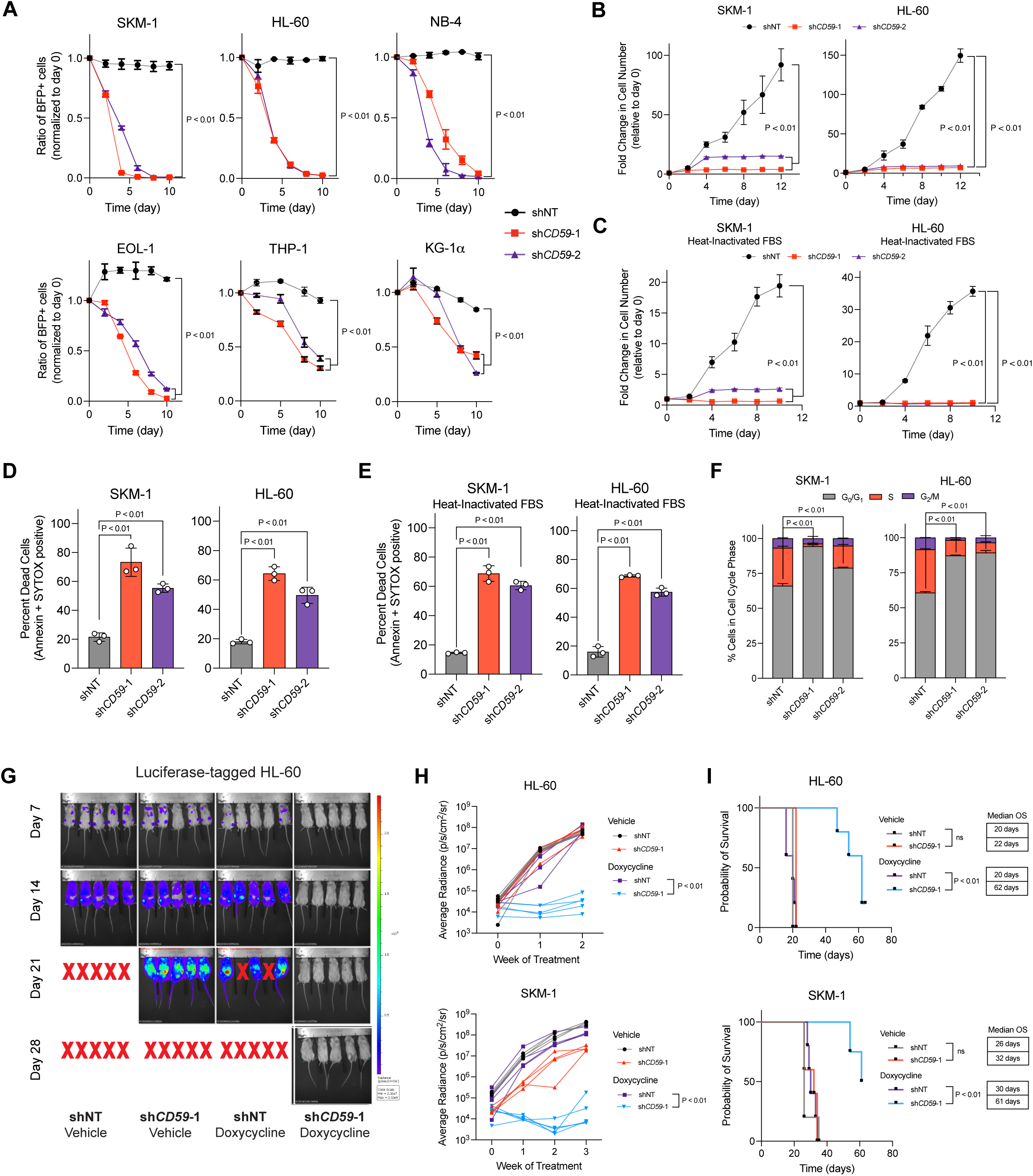
*CD59* silencing causes complement-independent cell-cycle arrest and impairs leukemic expansion in vivo. **(A)** Longitudinal competition experiment in six AML cell lines (SKM-1, HL-60, NB-4, EOL-1, THP-1, KG-1α) transduced with doxycycline-inducible sh*CD59*-1, sh*CD59*-2, or shNT lentiviral vectors. BFP^+^ and BFP^-^ cells were mixed at a 1:1 ratio and *CD59* knockdown was induced with 100 ng/ml doxycycline. The ratio of BFP^+^:BFP^-^ within each hairpin cells normalized to day 0 is shown over 10 days (p<0.01). **(B-C)** Fold change in cell number relative to day 0 in shNT, sh*CD59*-1, and sh*CD59*-2-expressing SKM-1 (left) and HL-60 (right) cells cultured in standard FBS (B) or heat-inactivated FBS (C) to exclude complement-mediated effects (p<0.01). **(D-E)** Cell death assessed by Annexin V and SYTOX staining at day 8 after doxycycline treatment in shNT, sh*CD59*-1, and sh*CD59*-2-expressing SKM-1 and HL-60 cells cultured in standard FBS (D) or heat-inactivated FBS (E) (p<0.01). **(F)** Cell-cycle analysis by DNA content and EdU incorporation in shNT, sh*CD59*-1-, and sh*CD59*-2-expressing SKM-1 (left) and HL-60 (right) cells. **(G)** Representative bioluminescence images of NSG mice transplanted with luciferase-tagged HL-60 cells expressing doxycycline-inducible shNT or sh*CD59*-1 and treated with vehicle or 60 mg/kg of doxycycline, shown at days 7, 14, 21, and 28. **(H)** Average radiance (p/s/cm²/sr) over weeks of treatment for HL-60 (top) or SKM-1 (bottom) CDX mice across all four treatment arms. **(I)** Kaplan-Meier overall survival curves for HL-60 (top) and SKM-1 (bottom) CDX mice in the vehicle and doxycycline treatment arms.

To confirm that the competitive disadvantage reflected loss of CD59, we purified BFP⁺ sh*CD59*- and shNT-expressing SKM-1 and HL-60 cells by cell sorting and verified CD59 downregulation at the mRNA and protein levels (**Extended Fig. 2E**). We next asked whether *CD59* depletion impairs growth outside a competitive setting, where cell-intrinsic effects could be obscured by the presence of untransduced cells. Cultured in isolation, sh*CD59*-expressing cells showed a marked reduction in absolute cell number relative to shNT controls in both lines (**Fig. 2B**), indicating that the requirement for CD59 is cell-intrinsic.

Because inhibition of MAC formation is the canonical function of CD59, we considered whether the growth impairment reflected complement activation^25^. This was unlikely as bovine complement has limited lytic activity against human cells^26^. Nonetheless, because these experiments were performed in medium containing non-heat-inactivated fetal bovine serum (FBS), we addressed the possibility directly by repeating them in medium supplemented with heat-inactivated FBS, which abrogates residual complement activity. *CD59* knockdown produced comparable reductions in cell number under these conditions (**Fig. 2C**), confirming that the effect is complement-independent.

We asked whether the reduced cell yield reflected increased cell death, decreased proliferation, or both. *CD59* knockdown increased cell death in both lines, and to a similar degree under standard and heat-inactivated conditions (**Fig. 2D, 2E**). In parallel, cell-cycle analysis of viable cells by DNA content staining and EdU incorporation revealed an accumulation of sh*CD59*-expressing cells in G0/G1 phase with a corresponding reduction in S phase, consistent with cell-cycle arrest (**Fig. 2F**). Together, these data identify a complement-independent role for CD59 in sustaining AML cell proliferation and survival.

Given the effects of *CD59* silencing on leukemic growth in vitro, we asked whether the in vivo microenvironment could compensate for loss of CD59. We therefore established luciferase-labeled cell line–derived xenograft (CDX) models to permit longitudinal monitoring of disease burden by bioluminescence imaging (BLI). NOD/SCID/IL2Rγ-null (NSG) mice were transplanted with sorted SKM-1 or HL-60 cells expressing doxycycline-inducible sh*CD59*-1 or shNT. Engraftment was confirmed by BLI on day 7, after which mice received daily oral doxycycline or vehicle until humane endpoint. After two weeks of treatment, doxycycline-treated mice transplanted with sh*CD59*-1-expressing cells showed an approximately 1,000-fold lower leukemic burden by BLI than the three control groups, comprising doxycycline-treated shNT recipients and vehicle-treated recipients of either construct (**Fig. 2G, H**). *CD59* knockdown also significantly prolonged survival relative to all control arms (**Fig. 2I**). Taken together, these findings demonstrate that CD59 is required for leukemic expansion in vivo.

### CD59 sustains MEK/ERK signaling by maintaining inner-leaflet phosphatidylserine and Ras/c-Raf membrane localization

To define the mechanisms by which CD59 supports AML proliferation and survival, we first performed bulk RNA sequencing of sh*CD59*-1 and shNT-expressing SKM-1 and HL-60 cells at 4 days after knockdown induction, a time point preceding appreciable cell death and therefore capturing the early transcriptional response to CD59 loss. Gene set enrichment analysis (GSEA) revealed downregulation of gene sets associated with cell proliferation, including E2F targets and S phase transition (**Fig. 3A; Extended Fig. 3A**), consistent with our cell cycle analysis (**Fig. 2F**). Because CD59 is a GPI-anchored surface protein lacking any cytoplasmic domain, these transcriptional changes must be relayed through intracellular signaling. To identify the responsible pathways, we performed unbiased phosphoproteomic profiling of the same cell models and applied the PhosR package to infer kinase activity from the phosphorylation status of annotated kinase substrates^27^. Activity of extracellular signal-regulated kinase 1 (ERK1), casein kinase 2 alpha (CK2α), and the cell cycle kinases CDK2 and CDK7 was predicted to be reduced in both cell lines following *CD59* silencing (**Fig. 3B**).

**Figure 3:**
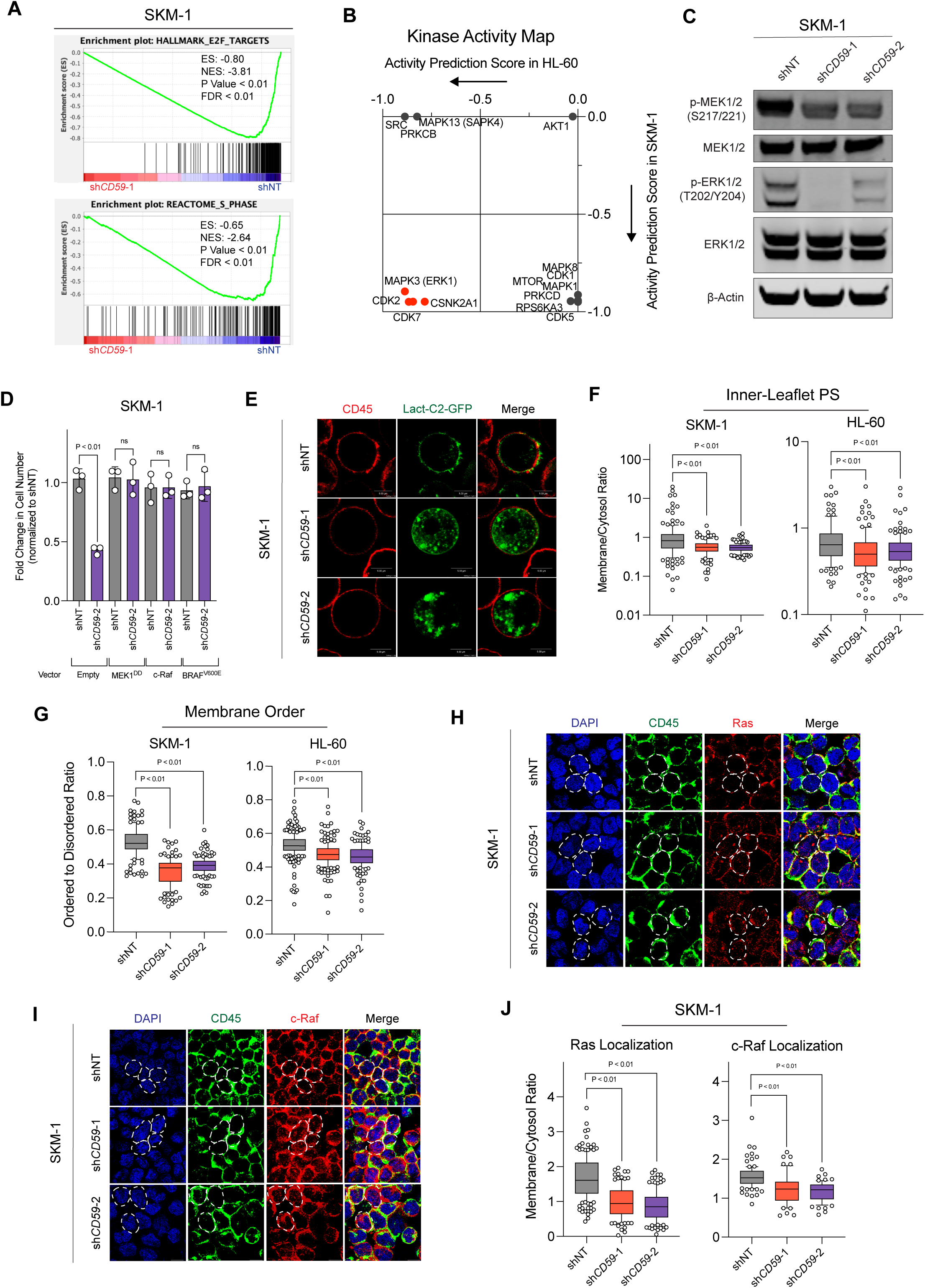
*CD59* silencing impairs ERK signaling through reduced inner leaflet phosphatidylserine and Ras/c-Raf membrane localization. **(A)** Gene set enrichment analysis (GSEA) of RNA-seq data from sh*CD59*-1 versus shNT-expressing SKM-1 cells. Representative enrichment plots for HALLMARK E2F targets (top) and REACTOME S PHASE targets (bottom). **(B)** Kinase activity map generated from phosphoproteomic profiling of sh*CD59*-1 and shNT-expressing SKM-1 and HL-60 cells using the phosR kinase-substrate prediction algorithm. ERK1, CK2α, CDK2, and CDK7 are among the most significantly reduced kinases in sh*CD59*-expressing cells across both cell lines. **(C)** Western blot for p-MEK1/2 (S217/221), MEK1/2, p-ERK1/2 (T202/Y204), ERK1/2, and β-actin in shNT-, sh*CD59*-1-, and sh*CD59*-2-expressing SKM-1 cells. **(D)** Fold change normalized to shNT-expressing cells for SKM-1 cells overexpressing empty vector (EV), constitutively active MEK1 (MEK1^DD^), wildtype c-Raf, or mutant BRAF (BRAF^V600E^). **(E)** Representative live cell images of shNT-, sh*CD59*-1, and sh*CD59*-2-expressing SKM-1 cells electroporated with Lact-C2-GFP (green), a phosphatidylserine inner-leaflet reporter, and co-stained with CD45 (red) as an outer membrane marker. White scale bars, 5 μm. **(F)** Quantification of membrane-to-cytosol ratio for inner-leaflet PS in shNT, sh*CD59*-1, and sh*CD59*-2-expressing SKM-1 and HL-60 cells, based on CD45-masked confocal image analysis (p<0.01). **(G)** Membrane order measured by Di-4-ANEPPDHQ staining in shNT, sh*CD59*-1, and sh*CD59*-2-expressing SKM-1 cells. **(H)** Representative confocal images of shNT, sh*CD59*-1, and sh*CD59*-2-expressing SKM-1 cells stained with DAPI (blue), CD45 (green), and anti-Ras antibody (red), showing reduced membrane Ras localization in sh*CD59*-expressing cells. White scale bars, 10 μm. **(I)** Representative confocal images of shNT, sh*CD59*-1, and sh*CD59*-2-expressing SKM-1 cells stained with DAPI (blue), CD45 (green), and anti-c-Raf antibody (red), showing reduced membrane c-Raf localization in sh*CD59*-expressing cells. White scale bars, 10 μm. **(J)** Quantification of membrane-to-cytosol ratio for Ras (left) and c-Raf (right) in shNT, sh*CD59*-1, and sh*CD59*-2-expressing SKM-1 cells, based on CD45-masked confocal image analysis (p<0.01).

We focused on the role of ERK1/2 signaling because it is constitutively activated in most AML patient samples and is well characterized to transmit proliferative and pro-survival signals^28,29^. Western blotting confirmed reduced phospho-ERK1/2 (T202/Y204) and reduced phosphorylation of the immediate upstream activator MEK1/2 (S217/S221), with total ERK1/2 and MEK1/2 unchanged (**Fig. 3C; Extended Fig. 3B**). To test whether suppression of MEK/ERK signaling is causal, we expressed a constitutively active MEK1 variant (MEK1^DD^) in sh*CD59*- and shNT-expressing SKM-1 cells and observed rescue of the growth impairment caused by *CD59* silencing (**Fig. 3D**). Together, these data indicate that CD59 sustains AML proliferation and survival at least in part by maintaining MEK/ERK signaling and its downstream transcriptional programs.

ERK1/2 are the terminal kinases of the Ras-Raf-MEK cascade, prompting us to ask where in this cascade CD59 loss interrupts signal transmission^30^. Both AML models carry activating RAS mutations, KRAS p.K117N in SKM-1 and NRAS p.Q61L in HL-60, which increase the GTP-bound fraction of Ras through intrinsic properties of the mutant proteins. Consistent with this, total Ras-GTP levels measured by pulldown with the Ras-binding domain (RBD) of c-Raf and total Ras and c-Raf levels were unchanged following *CD59* silencing (**Extended Fig. 3C, 3D**). This finding, together with the reduced MEK1/2 phosphorylation observed above (**Fig. 3C; Extended Fig. 3B**) and the rescue of growth by MEK1^DD^ (**Fig. 3D**), which bypasses c-Raf entirely, points to a defect in c-Raf activation. To test this directly, we asked whether restoring signal output at the level of c-Raf itself was sufficient to overcome CD59 loss. Overexpression of wild-type c-Raf or the constitutively active BRAF^V600E^ mutant restored the proliferation of sh*CD59*-expressing SKM-1 cells to that of shNT-expressing controls (**Fig. 3D**). Because BRAF^V600E^ signals to MEK independently of Ras-mediated recruitment and dimerization, its rescue places the defect upstream of MEK, while rescue by wild-type c-Raf indicates that c-Raf activation is the limiting step in CD59-depleted cells.

c-Raf activation is a membrane-dependent, multistep process. Ras-GTP recruits c-Raf to the plasma membrane through its RBD, after which the cysteine-rich domain (CRD) engages inner-leaflet phosphatidylserine (PS), releasing c-Raf from autoinhibition and stabilizing the complex for dimerization and activation^31–33^. PS is also required for Ras itself, most stringently for K-Ras, which is displaced from the plasma membrane when PS is depleted^34–36^. Across Ras isoforms, PS maintains the lateral segregation of the inner-leaflet lipid assemblies on which productive Ras nanoclustering and signal output depend^37^. Because GPI-anchored proteins are transbilayer coupled to inner-leaflet lipids, including PS, we asked whether CD59 loss alters PS distribution at the plasma membrane^38^. Using the inner-leaflet PS reporter Lact-C2-GFP, we found that PS redistributed away from the plasma membrane in sh*CD59*-expressing SKM-1 and HL-60 cells (**Fig. 3E-F, Extended Fig. 3E**). Annexin V staining showed no increase in outer-leaflet PS exposure, in contrast to staurosporine-treated controls, indicating PS mislocalization rather than a failure to maintain PS asymmetry (**Extended Fig. 3F**). Inner-leaflet PS also promotes ordered membrane domains, both by retaining cholesterol in the cytosolic leaflet and through cholesterol-dependent transbilayer coupling, so its loss would be predicted to reduce plasma membrane order^38–41^. Consistent with this, the polarity-sensitive dye Di-4-ANEPPDHQ, whose emission is blue shifted in liquid-ordered relative to liquid-disordered environments^42^, revealed a significant decrease in membrane order in sh*CD59*-expressing cells (**Fig. 3G**). To determine whether this membrane reorganization affects Ras and c-Raf localization, we quantified their membrane-to-cytosol ratio by immunofluorescence, using CD45 surface staining to delineate the plasma membrane. Both ratios were significantly reduced following *CD59* silencing in SKM-1 and HL-60 cells (**Fig. 3H-J; Extended Fig. 3G-I**), reflecting a redistribution of Ras and c-Raf away from the plasma membrane. Together, these findings support a model in which CD59 loss depletes inner-leaflet PS and alters plasma membrane order, thereby reducing coupling between Ras and c-Raf and interrupting signal transmission to ERK.

### CD59 is preferentially expressed in LSCs and is required for their self-renewal in vitro and in vivo

MEK/ERK signaling has been implicated in the maintenance of LSCs in murine models of AML^43^. Given our finding that CD59 sustains ERK signaling, we asked whether this dependency extends to LSCs. We first characterized CD59 surface expression across the leukemic hierarchy of OCI-AML-8973 and OCI-AML-8227, two patient-derived AML models in which LSC activity is functionally enriched within the CD34^+^CD38^-^ compartment^44^ (**Extended Fig. 4A**). In both models, CD59 surface expression was highest on the CD34^+^CD38^-^ LSC-enriched fraction relative to the more differentiated fractions (**Fig. 4A**). Consistent with this, proteomic profiling of primary AML samples showed higher CD59 protein in sorted cell fractions that engrafted in NSG mice (LSC+) than in those that did not (LSC-) (**Fig. 4B**), extending our earlier finding that *CD59* expression correlates with LSC gene signatures (**Extended Data Fig. 1C-D**).

**Figure 4:**
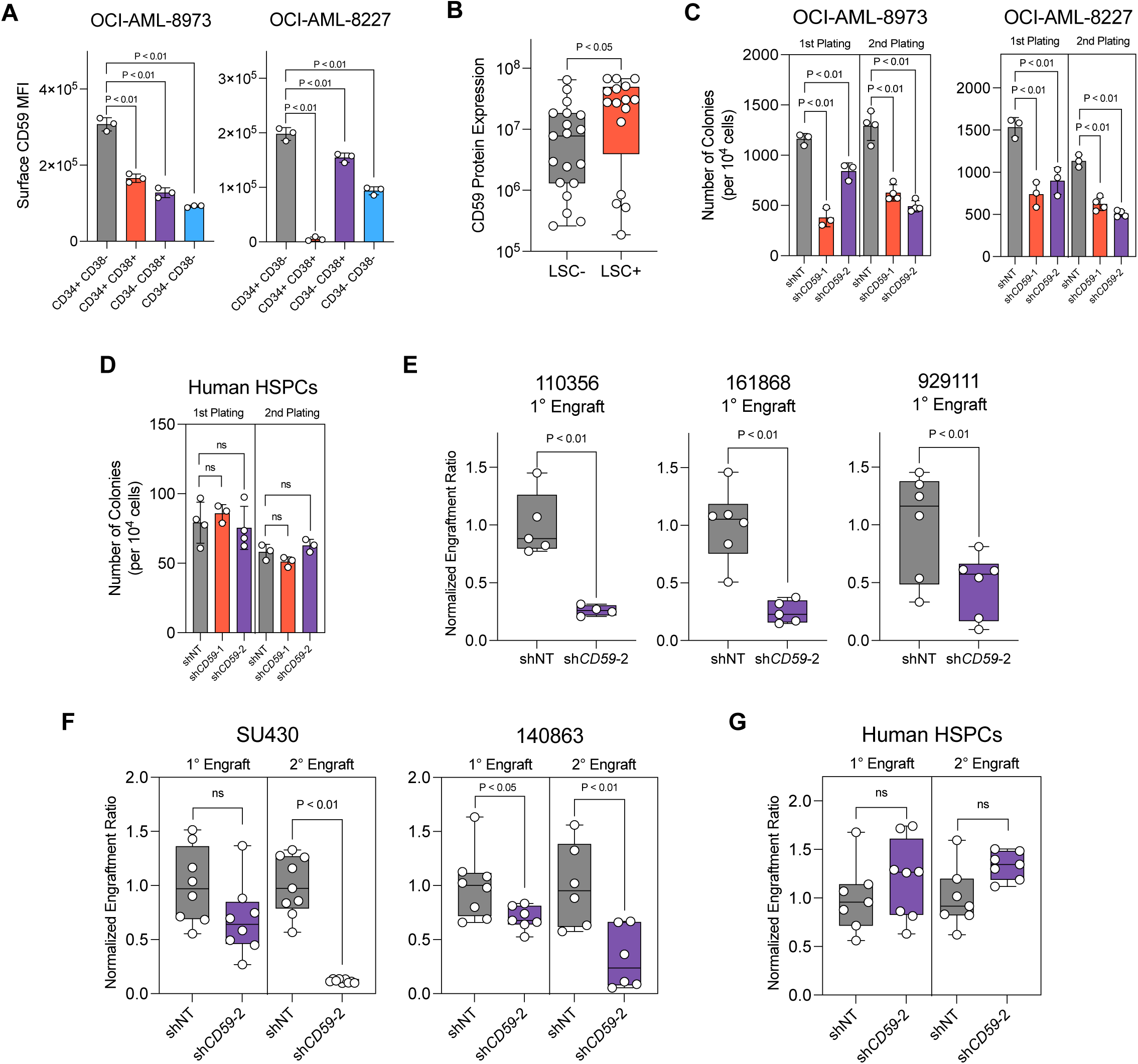
*CD59* downregulation impairs LSC self-renewal and reduces primary AML engraftment in vivo while sparing normal HSPCs. **(A)** Surface CD59 expression (MFI corrected to isotype control) across immunophenotypically defined compartments of the leukemic hierarchy in OCI-AML-8973 (left) and OCI-AML-8227 (right). **(B)** CD59 protein expression across LSC-non-engrafting and LSC+ engrafting fractions of primary AML samples. **(C)** Colony forming capacity of shNT, sh*CD59*-1, and sh*CD59*-2-expressing OCI-AML-8973 (left) and OCI-AML-8227 (right) cells across two serial platings. **(D)** Colony forming capacity of shNT, sh*CD59*-1, and sh*CD59*-2-expressing human HSPCs across two serial platings. **(E)** Normalized engraftment ratio of 110356, 161868, and 929111 primary AML cells transduced with shNT or sh*CD59*-2 in NSG mice. **(F)** Normalized engraftment ratio of SU430 and 140863 primary AML cells transduced with shNT or sh*CD59*-2 in primary and secondary NSG-recipients. **(G)** Normalized engraftment ratio of human HSPCs transduced with shNT or sh*CD59*-2 in primary and secondary NSG-recipients.

To test whether CD59 is functionally required in LSCs, we transduced both models with lentiviral vectors constitutively expressing sh*CD59*-1, sh*CD59*-2 or shNT together with a BFP reporter and confirmed knockdown of surface CD59 protein (**Extended Fig. 4B**). *CD59* knockdown significantly reduced colony-forming capacity across two serial platings in both models (**Fig. 4C**), indicating impaired clonogenic self-renewal. In contrast, *CD59* silencing did not impair the colony-forming capacity of CD34^+^ HSPCs isolated from human cord blood over two serial platings (**Fig. 4D; Extended Fig. 4C**), demonstrating selectivity for leukemic over normal clonogenic progenitors.

We next asked whether CD59 is required for the repopulating activity of primary human AML in vivo. Five patient AML samples were transduced with sh*CD59* or shNT vectors carrying a BFP reporter, and knockdown of CD59 protein was confirmed 24-48 hours post-transduction (**Extended Fig. 4D**). Unsorted transduced cells, comprising both BFP^+^ and BFP^-^ populations, were transplanted into sub-lethally irradiated NSG mice. Engraftment of human BFP^+^ cells was assessed by bone marrow aspiration 8 to 12 weeks post-transplant and expressed as the normalized engraftment ratio, defined as the representation of transduced cells in the xenograft normalized to input frequency and to shNT controls (**Extended Fig. 4E**). In three of the five samples, *CD59* silencing significantly reduced the relative engraftment potential in primary recipients (**Fig. 4E**). In the remaining two samples, the reduction in primary recipients was more modest, reaching significance in one sample but not the other (**Fig. 4F**). Since a defect restricted to the LSC compartment may not be apparent in primary recipients, we serially transplanted bone marrow cells from these recipients into secondary hosts. In both samples, *CD59* silencing significantly reduced the relative engraftment potential in secondary recipients (**Fig. 4F**), consistent with impaired LSC activity. In contrast, sh*CD59*-expressing CD34^+^ human HSPCs engrafted at levels comparable to shNT-expressing HSPCs in both primary and secondary recipients (**Fig. 4G**), indicating that CD59 is dispensable for normal human HSPC repopulation and supporting a therapeutic window for targeting CD59 in AML.

### rILYd4 depletes surface CD59 and selectively impairs AML proliferation

Our rationale for focusing on cell surface proteins was their accessibility to biologics, which confer a specificity that is difficult to achieve with small molecules. Because the case for targeting CD59 rests on a differential dependency rather than differential expression, we sought a biologic that blocks the ability of CD59 to sustain signaling, rather than one that delivers cytotoxicity. The mechanism described above predicts that depleting CD59 from the cell surface is a potential approach. This search led us to intermedilysin (ILY), a cytolytic toxin secreted by *Streptococcus intermedius* that binds human CD59 with high affinity and specificity through its fourth domain^45^. The 114 amino acid recombinant fourth domain, termed rILYd4, retains CD59 binding without inducing cell lysis and, importantly, triggers rapid lysosomal internalization and degradation of CD59 in cancer cells^46^. We therefore asked whether rILYd4 treatment recapitulates the effects of genetic knockdown in AML cells.

We generated and purified recombinant, polyhistidine (His)-tagged rILYd4, along with rILYd4^YYRSY^, a binding-deficient mutant harbouring alanine substitutions at key CD59-contact residues, as a control^47^ (**Extended Fig. 5A, 5B**). Incubating SKM-1 and HL-60 cells with the recombinant proteins at 4°C followed by anti-His-Tag staining and flow cytometric analysis showed a clear rightward shift with rILYd4 but not with rILYd4^YYRSY^, confirming the binding specificity of the wildtype protein for surface CD59 (**Extended Fig. 5C**). To determine whether rILYd4 binding causes CD59 degradation, we treated SKM-1 cells with rILYd4 or rILYd4^YYRSY^ for 8 hours at 37°C and immunoblotted whole cell lysates for total CD59. Treatment with rILYd4, but not rILYd4^YYRSY^, reduced CD59 protein levels relative to untreated controls (**Fig. 5A**). To study the mechanism, we performed confocal microscopy on treated SKM-1 cells. rILYd4 treatment drove internalization of CD59 and a reduction in CD59 staining, and the latter was prevented by the lysosomal inhibitor leupeptin (**Extended Fig. 5D**), consistent with lysosomal-mediated degradation.

**Figure 5.**
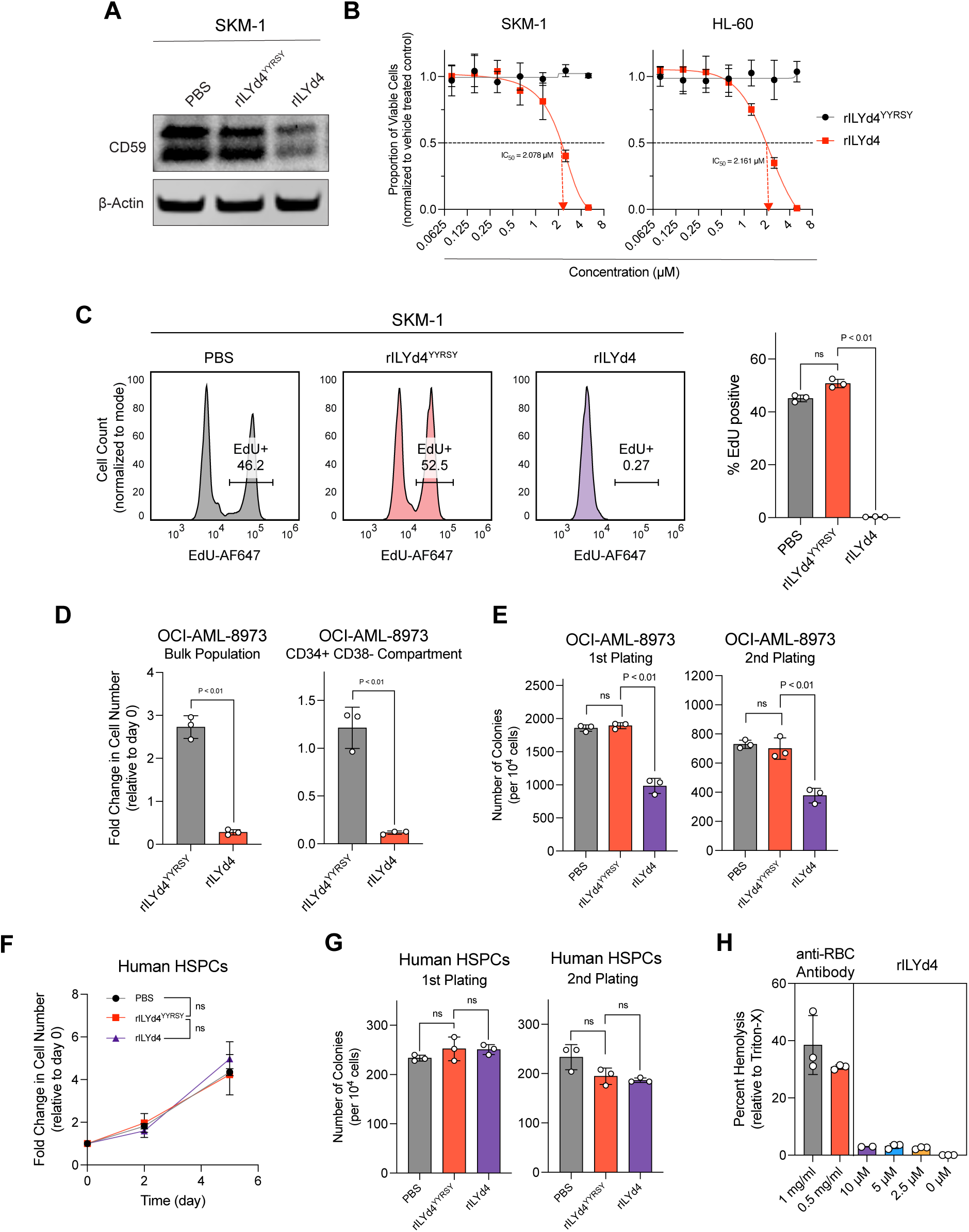
rILYd4 selectively depletes CD59 from leukemic cells, impairs LSC self-renewal, and does not impact human HSPCs. **(A)** Western blot for CD59 and β-actin in SKM-1 cells following 8-hour treatment with PBS, rILYd4^YYRSY^ or rILYd4. **(B)** Dose-response curves showing proportion of viable SKM-1 (left) and HL-60 (right) cells following treatment with increasing concentrations of rILYd4^YYRSY^ (black) or rILYd4 (red), normalized to vehicle-treated controls. IC_50_ values are indicated. **(C)** Cell-cycle analysis by EdU incorporation in PBS, rILYd4^YYRSY^, and rILYd4-treated SKM-1 cells. **(D)** Fold change in cell number in the bulk population (left) and CD34^+^CD38^-^ LSC-enriched compartment (right) of OCI-AML-8973 following treatment with rILYd4^YYRSY^ or rILYd4 (p<0.01). **(E)** Colony forming capacity of OCI-AML-8973 cells treated with PBS, rILYd4^YYRSY^, or rILYd4 at indicated concentrations across first (left) and second (right) platings (p<0.01). **(F)** Fold change in cell number relative to day 0 in CD34+ cord blood HSPCs treated with rILYd4^YYRSY^ or rILYd4 over 6 days. **(G)** Colony forming capacity of cord blood HSPCs treated with PBS, rILYd4^YYRSY^, or rILYd4 at indicated concentrations across first (left) and second (right) platings. **(H)** Hemolysis assay in isolated human red blood cells treated with anti-RBC antibody (positive control) or increasing concentrations of rILYd4 in the presence of 12.5% normal human serum.

We next asked whether depleting CD59 with rILYd4 phenocopies genetic knockdown. rILYd4 reduced the growth of SKM-1 and HL-60 cells with IC₅₀ values of 2.08 µM and 2.16 µM respectively (**Fig. 5B**), and abrogated EdU incorporation in both lines (**Fig. 5C; Extended Fig. 5E**). In contrast, rILYd4^YYRSY^ had no effect on the viability or proliferation of either cell line. Furthermore, rILYd4 reduced the expansion of not only the bulk leukemic population but also the CD34^+^CD38^-^ LSC-enriched fraction in OCI-AML-8973 and OCI-AML-8227 (**Fig. 5D; Extended Fig. 5F**), suggesting an impact on LSCs. Accordingly, rILYd4, but not rILYd4^YYRSY^, reduced the colony forming potential of OCI-AML-8973 cells across two platings, consistent with a reduction in LSC self-renewal (**Fig. 5E**). Together, these findings show that depletion of CD59 with rILYd4 recapitulates the effects of genetic knockdown, arresting leukemic proliferation and impairing LSC self-renewal.

To assess selectivity, we studied the effect of rILYd4 on normal hematopoietic cells. rILYd4 had no impact on the proliferation of CD34^+^ cord blood-derived HSPCs, nor on their colony forming capacity across two platings (**Fig. 5F, G**). Because CD59 protects erythrocytes from complement-mediated lysis, we also addressed the potential safety concern of hemolysis directly. Isolated human red blood cells treated with rILYd4 in the presence of 12.5% normal human serum showed minimal hemolysis at concentrations up to 10 µM, approximately five-fold above the concentration required to inhibit leukemic cell growth, whereas an anti-RBC antibody control produced a substantially higher degree of lysis (**Fig. 5H**). Collectively, these findings demonstrate that depletion of CD59 with rILYd4 impairs AML proliferation and LSC self-renewal while sparing normal hematopoietic progenitors and erythrocytes.

### rILYd4 reduces mitochondrial respiration and sensitizes AML cells to venetoclax

Venetoclax-based regimens have become standard of care for patients with AML who are unfit for intensive chemotherapy, and resistance to these regimens is a critical clinical problem^48,49^. Given that activation of the Ras-MAPK pathway has been shown to promote mitochondrial respiration and venetoclax resistance in AML cells^50^, we asked whether CD59 depletion lowers mitochondrial respiration and sensitizes AML cells to venetoclax.

Knockdown of *CD59* in SKM-1 and HL-60 cells significantly reduced both basal and maximal oxygen consumption rate (OCR) compared with shNT controls (**Fig. 6A**). rILYd4 treatment recapitulated this effect, indicating that targeted degradation of CD59 is also sufficient to impair mitochondrial respiration (**Fig. 6B, C**). Consistent with the link between mitochondrial OXPHOS and venetoclax sensitivity, sh*CD59*-expressing and rILYd4-treated SKM-1 and HL-60 cells showed significantly lower venetoclax IC_50_ values than controls (**Fig. 6D, E**), indicating that CD59 loss sensitizes AML cells to venetoclax.

**Figure 6.**
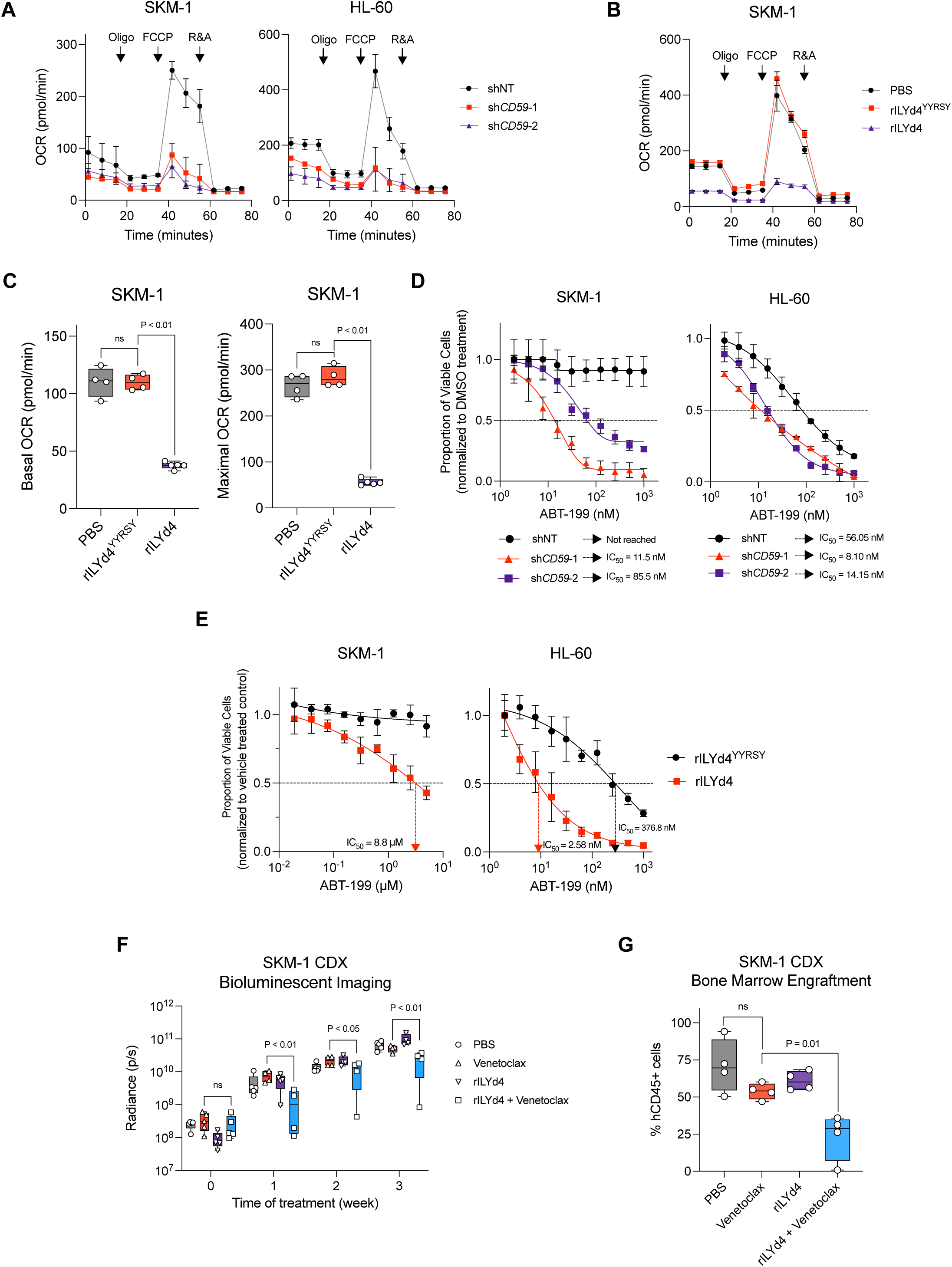
rILYd4-mediated CD59 degradation reduces mitochondrial respiration, sensitizes AML to venetoclax, and synergizes with venetoclax to reduce leukemic burden in vivo. **(A)** Mitochondrial stress test showing oxygen consumption rate (OCR) over time in response to oligomycin, FCCP, and rotenone/antimycin A (R&A) in shNT, sh*CD59*-1, and sh*CD59*-2-expressing SKM-1 (left) and HL-60 (right) cells **(B)** Mitochondrial stress test showing OCR in PBS, rILYd4^YYRSY^, and rILYd4-treated SKM-1 cells. **(C)** Basal OCR (left) and maximal OCR (right) measured by mitochondrial stress test in SKM-1 cells treated with PBS, rILYd4^YYRSY^, or rILYd4, demonstrating significantly reduced mitochondrial respiration following rILYd4 treatment (p<0.01). **(D)** Dose-response curves for venetoclax (ABT-199) in shNT, sh*CD59*-1, and sh*CD59*-2-expressing SKM-1 (left) and HL-60 (right) cells, showing significantly reduced IC_50_ values in sh*CD59*-expressing cells consistent with venetoclax sensitization. **(E)** Dose-response curves for venetoclax (ABT-199) in rILYd4^YYRSY^ and rILYd4-treated SKM-1 (left) and HL-60 (right) cells, showing increased venetoclax sensitization in the rILYd4-treated group. IC_50_ values are indicated. **(F)** Radiance (p/s) over weeks of treatment for SKM-1 CDX mice across all four treatment arms. **(G)** Bone marrow engraftment of SKM-1 cells from NSG mice treated with vehicle, venetoclax, rILYd4, and rILYd4 + venetoclax at endpoint.

Finally, we asked whether the impact of rILYd4 on venetoclax sensitivity could be reproduced in vivo. NSG mice were transplanted with luciferase-tagged SKM-1 cells which are intrinsically resistant to venetoclax (**Fig. 6D, E**). Seven days after transplantation, equivalent engraftment was confirmed by BLI before randomization to vehicle control, venetoclax (100 mg/kg/day by oral gavage), rILYd4 (10 mg/kg/day by intraperitoneal injection), or the combination, each given on a 5-days on, 2-days off schedule for 3 weeks. We selected an rILYd4 dose with limited single agent activity so that any sensitization to venetoclax would be apparent. Mice receiving the combination showed significantly lower leukemic burden by serial imaging (**Fig. 6F**) and, importantly, reduced human cell engraftment in the bone marrow at endpoint (**Fig. 6G**). Together, these findings indicate that rILYd4-mediated CD59 depletion remodels mitochondrial metabolism to sensitize AML cells to venetoclax.

## Discussion

In this study, we identify CD59 as a surface protein that is required for leukemic proliferation, survival, and LSC activity. High CD59 expression marks an adverse-risk, stem-like state and predicts poor prognosis in AML. CD59 is unique as a target antigen because it is defined by differential dependency rather than by differential expression relative to normal hematopoietic cells. We therefore focused on a biologic that triggers degradation of CD59 protein rather than one that kills antigen-positive cells. This agent, rILYd4, depletes surface CD59, phenocopies genetic knockdown, and sensitizes AML cells to venetoclax. Together, these findings establish a surface target defined by functional dependency and a biologic that acts by lowering its abundance.

CD59 is best characterized as an inhibitor of the terminal complement pathway, yet our data reveal a complement-independent function in sustaining Ras-MAPK signaling. Non-canonical signaling roles have been described previously, including activation of lipid raft-associated tyrosine kinases, JAK2, and T cell receptor proximal signaling, but most relied on antibody-mediated crosslinking of CD59, and the physiological relevance of forced clustering is uncertain^51,52^. We show instead that CD59 governs Ras and c-Raf coupling by maintaining inner-leaflet PS, which is required for their activation and recruitment to the plasma membrane. This is perhaps unsurprising given that GPI-anchored proteins can couple the outer and inner leaflets, although the precise mechanism by which CD59 exerts this effect will require further study. Notably, a prior study also linked CD59 to Ras compartmentalization but reached an opposite functional conclusion^53^. In T cells, an intracellular pool of CD59 promoted Ras trafficking to the plasma membrane, and CD59 deficiency arrested Ras in the Golgi, thereby enhancing Ras-MAPK signaling. Both studies therefore indicate that CD59 regulates Ras membrane localization but differ in the downstream consequence, which may reflect the signaling competence of Golgi-localized Ras in lymphocytes as well as differences in membrane lipid composition and Raf isoform usage between AML blasts and T cells. Whether surface and intracellular CD59 pools have distinct functions remains an important unresolved question.

An important consideration for targeting CD59 is its therapeutic window. Given the role of CD59 in restraining complement-mediated lysis, hemolysis is the principal anticipated risk, as indicated by the phenotype of *Cd59a/b* knockout mice^17^. However, rILYd4 alone triggers minimal hemolysis in vitro, which we confirmed in human erythrocytes exposed to normal human serum at concentrations that suppressed AML growth. Furthermore, a prior study showed that one month of systemic rILYd4 administration caused no hemolysis or overt toxicity in human CD59 transgenic mice on a murine *Cd59*-deficient background^54^. This lack of toxicities likely reflects partial rather than complete CD59 inhibition together with redundancy from other complement regulatory proteins. Combined with preserved HSPC engraftment after *CD59* knockdown, these findings suggest a workable window, although formal toxicity studies in humanized models will be essential, since rILYd4 binds only human CD59. Immunogenicity is a second consideration for a bacterially derived biologic but is likely not prohibitive, as the long-standing clinical use of asparaginase for the treatment acute lymphoblastic leukemia illustrates, and PEGylation or epitope deimmunization could mitigate anti-drug responses while extending half-life. Further engineering to improve potency and add leukemia-directed targeting would potentially widen the therapeutic window further.

Clinically, the most immediate opportunity for CD59 degradation lies in combination therapy for adverse-risk AML, where venetoclax-based regimens are limited by intrinsic and acquired resistance^55^. Our data position CD59 degradation as a rational partner for BCL-2 inhibition rather than as a single agent, with CD59-high and RAS-MAPK-activated disease the most rational starting point. This strategy may also have broader applicability. High CD59 expression predicts adverse outcomes across multiple solid tumors, an association usually attributed to complement resistance, but a cell-intrinsic role in sustaining Ras-MAPK signaling offers an alternative explanation^38,56–60^. Targeted degradation of CD59 may therefore be worth evaluating in other cancer types.

Several limitations warrant emphasis. Our studies using primary AML samples were limited in number, and larger cohorts spanning defined genetic subgroups will be needed to establish whether specific mutational profiles predict CD59 dependency. Relatedly, why adverse-risk genetics are associated with higher CD59 expression is unknown, and defining the transcriptional or epigenetic basis of this relationship warrants further investigation. Our combination data are restricted to venetoclax, and systematic screening for additional partners is warranted. Finally, how CD59 controls inner-leaflet PS remains unclear and will require dedicated lipid biochemistry to resolve.

In summary, CD59 sustains Ras-MAPK signaling through plasma membrane organization, is preferentially required by LSCs, promotes venetoclax resistance, and can be depleted by rILYd4, a targeted biologic that spares normal progenitors. These findings establish CD59 as an actionable vulnerability and provide proof of concept its targeted degradation is a viable therapeutic modality in AML.

## Materials and Methods

### Tissue culture

SKM-1, HL-60, EOL-1, NB-4, THP-1, KG-1a, MOLM-13, OCI-AML-3, and NOMO-1 cells were cultured in RPMI-1640 medium (Wisent Bioproducts, #350-000) with 20% (v/v) fetal bovine serum (FBS) (Wisent Bioproducts, #080-150), and 1% (v/v) GlutaMax (Gibco, #35050061). All cell lines were routinely tested for mycoplasma using an ELISA-based detection kit and periodic STR-profiling was performed to ensure cell-line identity.

OCI-AML-8973, OCI-AML-8227, and primary AML samples were cultured in X-VIVO® 15 Serum-free Hematopoietic Cell Medium (Lonza, #02-060Q) supplemented with 20% (v/v) BIT 9500 Serum Substitute (STEMCELL, #09500), 1% (v/v) GlutaMax (Gibco, #35050061), and supplemented with the following cytokines: 10 ng/mL human recombinant IL-6 (STEMCELL, #78050), 10 ng/mL human recombinant IL-3 (STEMCELL, #78040), 50 ng/mL mouse recombinant SCF (STEMCELL, # 78064), 50 ng/mL, human recombinant Flt3 Ligand (STEMCELL, # 78009), 10 ng/mL human recombinant G-CSF (STEMCELL, # 78012), and 25 ng/mL human recombinant TPO (STEMCELL, # 78210.1).

Human cord blood samples were cultured in StemSpan™-XF (STEMCELL, #100-0073) supplemented with 2% (v/v) GlutaMax, 100 ng/mL of mSCF, 100 ng/mL of hFlt3 ligand, 20 ng/mL of hTPO, 1 µM of SR1 (STEMCELL, #72342), and 50 nM of UM171 (STEMCELL, #72912).

### *In vitro* competition assays

AML cell lines transduced with shNT or sh*CD59*-1/2-expressing lentiviral vectors were evaluated for stable BFP expression 4-days post-transduction. If the proportion of BFP^+^ cells were > 50% at baseline, untransduced cells were added to the culture to obtain a baseline BFP^+^ expression of ∼ 50%.

1 × 10^4^ cells were seeded in a 96-well flat-bottom plate in RPMI-1640 supplemented with 20% (v/v) FBS, 1% (v/v) GlutaMax, and 100 ng/ml of doxycycline (MedChem Express, #24390-14-5). The proportion of BFP+ cells was evaluated every 2 days using a BECKMAN CytoFlex flow cytometer. Doxycycline was replenished at each media change every 2 days.

### Cell-line derived xenografts

SKM-1 and HL-60 cells were transduced with pRSITE-sh*CD59*-1-BFP or pRSITE-shNT-BFP and pHIV-Luc-ZsGreen. BFP^+^ and GFP^+^ double positive cells were sorted via fluorescence activated cell sorting (FACS) and 1 × 10^6^ cells were transplanted into 6–8-week-old female NSG mice conditioned with 250 rads of irradiation. Baseline engraftment was confirmed 7 days post-transplant using bioluminescence imaging by intraperitoneal injection of luciferin at a final dose of 150 mg/kg.

sh*CD59*-1 or shNT-expressing cells were either treated with water or 60 mg/kg of doxycycline via oral gavage and bioluminescence imaging was performed every 7 days to measure tumor burden until the mice reached endpoint.

### *In vivo* competition experiments

Human AML samples from peripheral blood or bone marrow biopsies were obtained with written consent according to procedures approved by the University Health Network ethics committee. Freshly thawed primary AML samples were subjected to T cell depletion using the EasySep™ Human CD3 Positive Selection Kit II (STEMCELL, #17851). Primary AML or cord blood cells were transduced with pRSI9-sh*CD59*-1/2-BFP or pRSI9-shNT-BFP and baseline level of BFP expression was measured 24-hours post-transduction. 1-2 × 10^6^ T cell-depleted primary AML cells or 1 × 10^5^ cord blood HSPCs were resuspended in 100 uL of Opti-MEM (ThermoFisher Scientific, #31985-070) and transplanted via tail vein injection into 6–8-week-old female NSG mice conditioned with 250 rads of irradiation. Bone marrow aspiration was performed on the left femur 8-12 weeks following transplantation. Red blood cell lysis was performed using RBC lysis buffer (BioLegend, #420301) for 5 minutes at room temperature and the level of hCD45-APC^+^ mCD45.1-APCFire750^-^ TER119-FITC^-^ Helix-Green^-^ BFP^+^ cells was measured.

For secondary engraftment, bone marrow cells from primary recipient mice were collected at endpoint. Red blood cell lysis was performed, and the samples were pooled and BFP+ expression was recorded. An equivalent number of cells were resuspended in 100 uL of Opti-MEM and transplanted into NSG mice conditioned with 250 rads of irradiation. Bone marrow samples were collected at 4-8 weeks following transplantation and the level of hCD45-APC^+^ mCD45.1-APCFire750^-^ TER119-FITC^-^ Helix-Green^-^ BFP^+^ cells was measured.

### Calculation of normalized engraftment ratio

We recorded the percentage of BFP^+^ live cells 24 hours after transduction to establish the baseline percentage of injected shNT and sh*CD59*-expressing cells. After engraftment, we measured the percentage of BFP^+^ cells within the human CD45^+^mCD45.1^−^ gate to determine the percentage of shRNA-expressing cells that had engrafted. For each sample, we divided the engrafted percentage of BFP^+^ cells by the baseline percentage of BFP^+^ cells to obtain a baseline-normalized engraftment value. We then divided each of these values by the mean baseline-normalized engraftment value of shNT-expressing cells within the same primary sample. The resulting scores reflect the normalized engraftment ratio of sh*CD59*-expressing cells relative to shNT controls.

### Human cord blood samples

Human cord blood samples were obtained with informed consent from Princess Margaret Cancer Centre following the procedures approved by the University Health Network (UHN) research ethics board. Mononuclear cell isolation and HSPC enrichment were conducted using Ficoll Paque Plus (Sigma Aldrich, #GE17-1440-02) and Human Progenitor Cell Enrichment Kit (STEMCELL, #19356).

### Lentivirus production and transduction

HEK293T cells were grown in DMEM (Wisent, #319-005) supplemented with 10% (v/v) FBS (Wisent, #080-150). Cells were seeded in 15 cm tissue culture plates at a density of 7 × 10^6^ cells per plate one day before transduction. The following day, cells were transfected with lentiviral plasmid vectors, psPAX2 (Addgene, #12259) and pCMV-VSVG (Cell biolabs, #320023) using the jetPRIME transfection reagent (Polyplus, #CA89129-924) according to the manufacturer’s protocol. Supernatant containing lentiviral particles was collected at 48-and 72-hours post transfection and filtered through a 0.22 µm PVDF membrane. Viral particles were precipitated in 40% (w/v) polyethylene glycol (Sigma, #89510-1KG-F) overnight. The viral pellet was collected by centrifugation at 3,148 × g for 30 minutes at 4 °C. The pellet was resuspended in HBSS (Gibco, #14170112) + 25 mM HEPES (ThermoFisher, #15630-080) and stored at −80°C for long term storage.

For lentiviral transductions, non-TC-treated 24 well plates were coated with 25 µg/mL of Retronectin (Takara, Cat # T100B) for 2 hours at room temperature followed by aspiration and blocking with PBS containing 2% (w/v) BSA (Wisent Bioproducts, Cat # 800-096-EG) for 30 minutes at room temperature. After aspiration of the blocking buffer, the concentrated virus suspension was added to wells. The plates were then centrifuged at 3,248 × g for two hours at 4°C to allow virus binding. Following centrifugation, unbound virus was aspirated, and cells were added. The plates were then transferred to a 37 °C incubator to initiate lentiviral infection.

### rILYd4 expression and purification

A pET30-rILYd4 plasmid was obtained from Biomatik. BL21(DE3+) *E. coli* cells were transformed with pET30-rILYd4 and plated on LB agar containing 50 µg/mL kanamycin. The following day, a starter culture was inoculated by resuspending a single colony in a 50 mL LB Broth culture overnight. The following morning, the starter culture was transferred to a 2 L culture of LB Broth and expression of rILYd4 was induced with 0.25 mM of IPTG once the culture reached an OD of ∼0.4. The protein was expressed overnight at 16°C. On the next day, the cells were centrifuged at 8,500 × g for 10 minutes and lysed with 50 mM Tris, 250 mM NaCl, 20 mM imidazole, 10% (v/v) glycerol, protease inhibitor, 1 mM PMSF, 10 µg/mL DNase I, and 100 mg/mL lysozyme. The solution was rotated at 4°C for 30 minutes and sonicated for 10 minutes. The resulting solution was ultracentrifuged at 40,000 × g for 45 minutes at 4°C and the supernatant was applied to His GraviTrap™ columns (Cytvia, #11003399). The protein was eluted with 250 mM of imidazole in the lysis buffer.

Purity was evaluated to be >90% via SDS-PAGE analysis and Coomassie blue staining, the peptide was dialyzed overnight to remove imidazole, and endotoxin removal was performed using Pierce™ High Capacity Endotoxin Removal Spin Columns (Thermo Scientific, #88274).

### Confocal microscopy

Standard cell mounting and imaging protocol was used. 2 × 10^5^ cells were resuspended in 100 µL of PBS and spun down onto a glass microscope slide using a Cytoclip Slide Clip (Fischer Scientific, #59910052) and reusable sample chamber on a Cytospin 4 centrifuge (Fischer Scientific, #29618304). The cells were fixed using 4% paraformaldehyde at room temperature for 15 minutes. The reaction was quenched using 0.1 M Glycine and the cells were permeabilized with 0.05% saponin. The cells were blocked in 5% (w/v) BSA, 1% Tween-20 and 0.05% saponin in PBS. The cells were incubated in a humidified chamber with primary antibodies overnight at 4°C in blocking buffer. The following morning, the cells were washed 3 times with the blocking buffer and stained for 1 hour at RT with the following secondary antibodies: Anti-rabbit IgG (H+L) Alexa Fluor 488 (CST, # 4412) and Anti-mouse IgG (H+L) Alexa Fluor 647 (CST, #4410). Finally, the cells were washed 3 times with blocking buffer and stained with ProLong Gold Antifade Reagent with DAPI (CST, #8961) and allowed to cure overnight at room temperature.

The slides were imaged using a Leica SP8 confocal microscope at the Advanced Optical Microscopy Facility at Princess Margaret Cancer Research Tower.

### Live cell imaging

For Lact-C2-GFP imaging, SKM-1 and HL-60 cells were nucleofected with Lact-C2-GFP using a Lonza nucleofector. The next day, the cells were stained with anti-human CD45-APC for 30 minutes at 4°C. For membrane order analysis, SKM-1 and HL-60 cells were stained with 5 µM of Di-4-ANEPPDHQ for 2 hours and washed 3 times with PBS. Next, the cells were transferred to Nunc™ Lab-Tek™ Chamber Slide System (Fischer Scientific, #1256522) and imaged using a Leica Stellaris live cell microscope. Cells were maintained in 5% CO_2_ at 37°C during all live cell imaging sessions.

### Ras/c-Raf membrane to cytosol quantification

Ras and Raf membrane localization was quantified from confocal images using CD45 as a plasma membrane boundary marker. Single-channel CD45 and Ras or c-Raf TIFF images were analyzed in Python. CD45 images were used to segment individual cells using Cellpose, and small debris or poorly segmented objects were excluded by area filtering. For each segmented cell, a membrane-proximal region was defined as the outer 2–3 pixels of the CD45-derived cell mask, while the cytosolic region was defined by eroding the cell mask inward and excluding membrane-adjacent pixels. Ras or Raf fluorescence intensity was measured in each compartment, and membrane enrichment was calculated as the mean membrane-proximal intensity divided by the mean cytosolic intensity. Measurements were performed at the single-cell level and summarized for statistical comparison between shNT and *CD59* knockdown conditions.

The python code can be found at: https://github.com/abdulamaher/CD59-in-AML/.

### Di-4-ANEPPDHQ membrane order analysis

Di-4-ANEPPDHQ membrane order images were quantified using a custom Python pipeline. For each field, ordered- and disordered-emission channel images were imported and analyzed together with Cellpose-derived cell masks. Individual segmented objects were treated as single cells, and per-cell regions of interest were used to extract fluorescence intensities from both Di-4 channels. Pixels were retained for analysis only if they passed total Di-4 intensity filtering, defined from the summed ordered and disordered channel intensities, and if they did not contain saturated or invalid values. For each valid pixel, membrane order was calculated as the mean ordered intensity divided by the mean disordered intensity.

The python code can be found at: https://github.com/abdulamaher/CD59-in-AML/.

### Red blood cell hemolysis assay

To isolate red blood cells, 15 mL of venous blood was collected from healthy donors using sterile technique in blood collection tubes coated with EDTA. Blood was centrifuged for 5 minutes at 400 × g to collected red blood cells which were washed 3 times in PBS. 10 μL of the RBC packed suspension was diluted in 990 μL of PBS to obtain a 1% RBC cell suspension.

Normal human serum (NHS) was isolated from venous blood of healthy donors as a source of complement in sterile blood collection tubes without anti-coagulants. The blood was allowed to clot at room temperature for 30 minutes following which the tubes containing the blood were centrifuged at 1000 × g for 10 minutes at room temperature. The supernatant containing the serum was transferred into a sterile 15 mL conical tube and placed on ice to preserve complement activity.

50 μL of 1% RBC cell suspension was added to 100 μL of GVB++ complement buffer (CompTech, #B102) containing anti-RBC complement activating antibody, rILYd4, or PBS and NHS at a final concentration of 12.5% (v/v). As a negative control, NHS was heat-inactivated at 56°C for 30 minutes and 1% (v/v) Triton-X was used as a positive control. The cell suspension was incubated at 37°C for 1 hour and centrifuged at 400 × g for 5 minutes at RT. 100 μL of the supernatant was transferred to a new 96-well flat-bottom plate and hemoglobin release was measured at 415 nm using a plate reader. Hemolysis was calculated using the following formula:

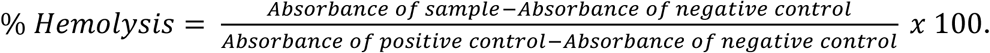

### EdU and cell cycle analysis

One day prior to the experiment, SKM-1 and HL-60 cells were seeded in a 24-well tissue culture plate at a cell density of 2.5 × 10^5^ cells/mL. The following day, EdU dissolved in DMSO was directly added to the growth medium at a final concentration of 10 μM. The cells were labelled with EdU for 1 hour at 37 °C. EdU staining was performed using the Click-iT™ EdU Alexa Fluor™ 647 Flow Cytometry Assay Kit (Invitrogen, #C10419) as per the manufacturer’s protocol.

### Colony-forming unit (CFU) assays

CD34^+^ human HSPCs were enriched from cord blood samples using the CD34^+^ positive selection kit (STEMCELL, #17856). 1 × 10^4^ CD34^+^ human HSPCs were resuspended in 1.1 mL of MethoCult™ H4434. The resuspended cells were treated with the relevant compounds (PBS, rILYd4^YYRSY^, or rILYD4) or plated as is to 35 mm cell culture dishes for HSPCs expressing sh*CD59* or shNT-targeting shRNAs. The number of colonies were counted after 10 days in culture, and the cells were replated if necessary.

### RNA-sequencing analysis

Total RNA was extracted from SKM-1 and HL-60 cells using the EZ-10 Spin Columns Total RNA Miniprep Kit (BioBasic, #BS1361) and quantified using a Nanodrop spectrophotometer. All samples were quality checked at the Princess Margaret Genomics Centre and had a RIN greater than 9. Samples were submitted to Novogene Corporation for sequencing analysis. Reads were mapped to reference genome using STAR software v2.6.1d.

For differential gene expression analysis, triplicate pairwise samples were analyzed using DESeq2 R package (v1.20.0). P-values were adjusted using the Benjamini and Hochberg’s approach. |log2FC| > 1.5 and P-Adjusted < 0.01 were used as cutoffs for differentially expressed genes.

### Phosphoproteomic profiling analysis

A total of 2.5 × 10^7^ SKM-1 or HL-60 cells expressing shNT or sh*CD59*-1 were collected 4 days after doxycycline induction in biological triplicates. Cell pellets were flash-frozen in liquid nitrogen and submitted to the SPARC BioCentre Molecular Analysis Centre at The Hospital for Sick Children, Toronto, ON, for phosphoproteomic analysis. Following cell lysis and protein digestion, samples underwent phosphopeptide enrichment targeting phosphorylated serine, threonine, and tyrosine residues. Enriched samples were analyzed by data-independent acquisition mass spectrometry. Raw mass spectrometry data were searched against the UniProt human Swiss-Prot database, including isoforms. Phosphosite abundance values were subsequently processed and analyzed using the phosR package.

### Active-RAS (GTP-Ras) pull-down assay

Active, GTP-bound Ras was isolated using the Active Ras Detection Kit (Cell Signaling Technology, #8821). shNT, sh*CD59*-1, and sh*CD59*-2-expressing SKM-1 and HL-60 cells were lysed in ice-cold 1X Lysis Buffer with 1 mM PMSF; lysates were clarified at 16,000 × g for 15 min at 4 °C and quantified by BCA assay. For each pull-down, ≥500 µg of lysate was incubated with 80 µg GST-Raf1-RBD and glutathione resin for 1 h at 4 °C, washed three times, and eluted in 2X reducing SDS sample buffer (200 mM DTT). Where indicated, lysates were pre-loaded with 0.1 mM GTPγS (positive control) or 1 mM GDP (negative control) in the presence of 10 mM EDTA prior to pull-down. Eluates were probed with Ras Monoclonal Antibody #8832 (1:200), which detects H, K, and N Ras using standard Western blotting techniques.

### Western blot

Standard Western blotting techniques were performed. Blots were incubated with primary antibodies diluted in 5% (w/v) BSA in TBST overnight at 4 °C on a shaking platform. The following next day, blots were washed for 5 minutes for three times with TBST and incubated with a secondary antibody (LI-COR, #926-32213 or LICOR, #926-68073) at 1:15,000 dilution in 5% (w/v) BSA in TBST at room temperature for 1 hour. Membranes were washed three times again and imaged on the Odyssey CLx Imaging system (LI-COR Biosciences).

### RT-qPCR

Total RNA was extracted from cells using the EZ-10 Spin Columns Total RNA Miniprep Kit (BioBasic, #BS1361) and quantified on a Nanodrop spectrophotometer. Reverse transcription and quantitative PCR were performed using the Luna® Universal One-step RT-qPCR kit (NEB, #E3005S) and the Bio-Rad CFX touch real-time PCR detection system. The threshold cycle (Ct) value was determined using CFX Manager v3.1. Gene expression was calculated using the ΔΔCt method. Ct values were normalized to beta-actin.

---

**Extended Figure 1:**
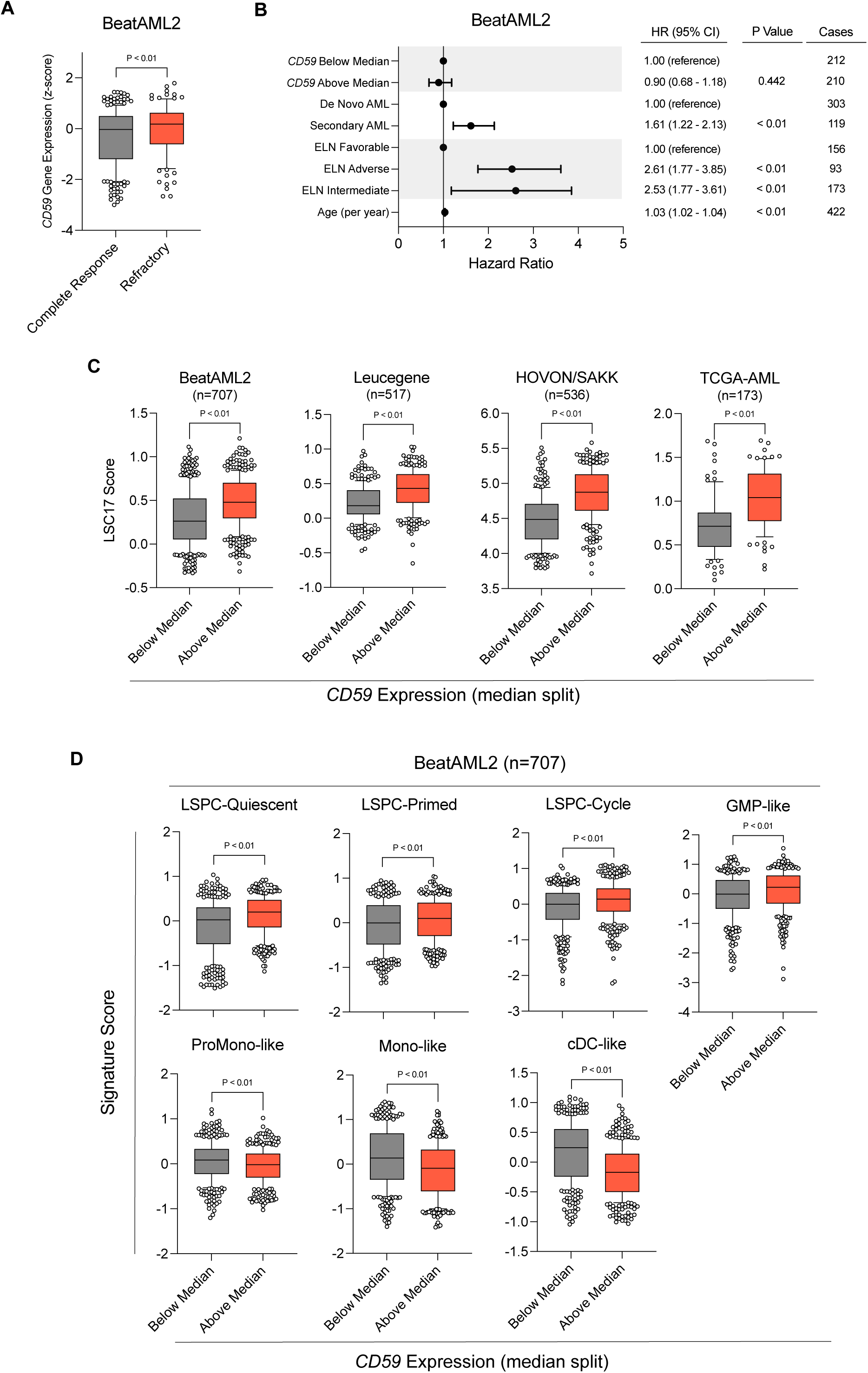
*CD59* expression associates with refractory disease, adverse-risk features, and stem/progenitor transcriptional programs in AML. **(A)** *CD59* expression (z-score) in BeatAML2 patients achieving a complete response versus those with primary refractory disease. Boxes show median and interquartile range, points are individual patients. **(B)** Multivariable Cox proportional-hazards analysis of overall survival in BeatAML2 (n = 422), modeling *CD59* expression (above vs below median), AML ontogeny (secondary vs de novo), ELN2017 risk (Adverse and Intermediate vs Favorable), and age (per year). Points and horizontal bars are hazard ratios and 95% confidence intervals; the table lists HR (95% CI), P value (Wald test), and case numbers per stratum. Reference categories are indicated. **(C)** LSC17 stem-cell score in patients stratified by *CD59* expression above versus below the cohort median, across four independent AML cohorts: BeatAML2 (n = 707), Leucegene (n = 517), HOVON/SAKK (n = 536), and TCGA-AML (n = 173). Boxes show median and interquartile range, points are individual patients. **(D)** Cell-state signature scores across the leukemic hierarchy in BeatAML2 (n = 707), stratified by CD59 expression above versus below median. Scores for stem/progenitor states (LSPC-Quiescent, LSPC-Primed, LSPC-Cycle, GMP-like) and differentiated states (ProMono-like, Mono-like, cDC-like) are shown; each signature score is the mean per-gene z-score of its member genes. Boxes show median and interquartile range, points are individual patients.

**Extended Figure 2.**
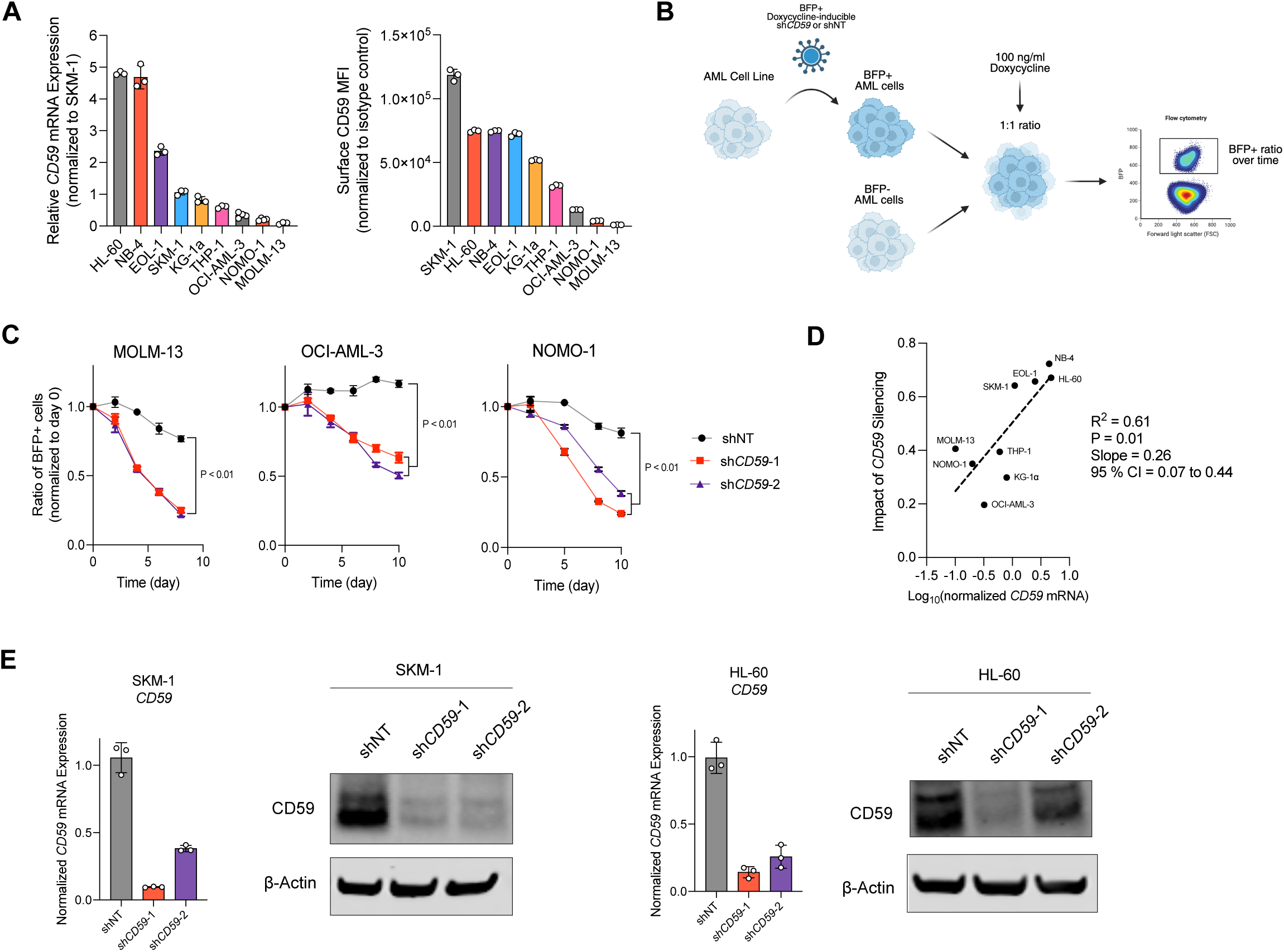
Characterization of CD59 expression and shRNA-mediated knockdown across AML cell lines. **(A)** *CD59* mRNA expression (left, normalized to SKM-1) and surface CD59 protein expression by flow cytometry (right, corrected to isotype control) across a panel of nine AML cell lines. **(B)** Schematic of the competitive growth assay. AML cell lines were transduced with doxycycline-inducible BFP-tagged shCD59 or shNT lentiviral vectors. BFP^+^ and BFP^-^ cells were mixed at a 1:1 ratio, treated with 100 ng/ml doxycycline, and BFP^+^ cell ratio was tracked longitudinally by flow cytometry. **(C)** Longitudinal competitive fitness assay in MOLM-13, OCI-AML-3, and NOMO-1 cell lines. **(D)** Correlation between baseline *CD59* mRNA expression and the magnitude of growth impairment following *CD59* silencing across nine AML cell lines. **(E)** Validation of *CD59* knockdown in FACS-isolated BFP^+^ shNT, sh*CD59*-1, and sh*CD59*-2-expressing SKM-1 and HL-60 cells at the mRNA level (bar graphs) and protein level (western blot), with β-actin as loading control.

**Extended Figure 3:**
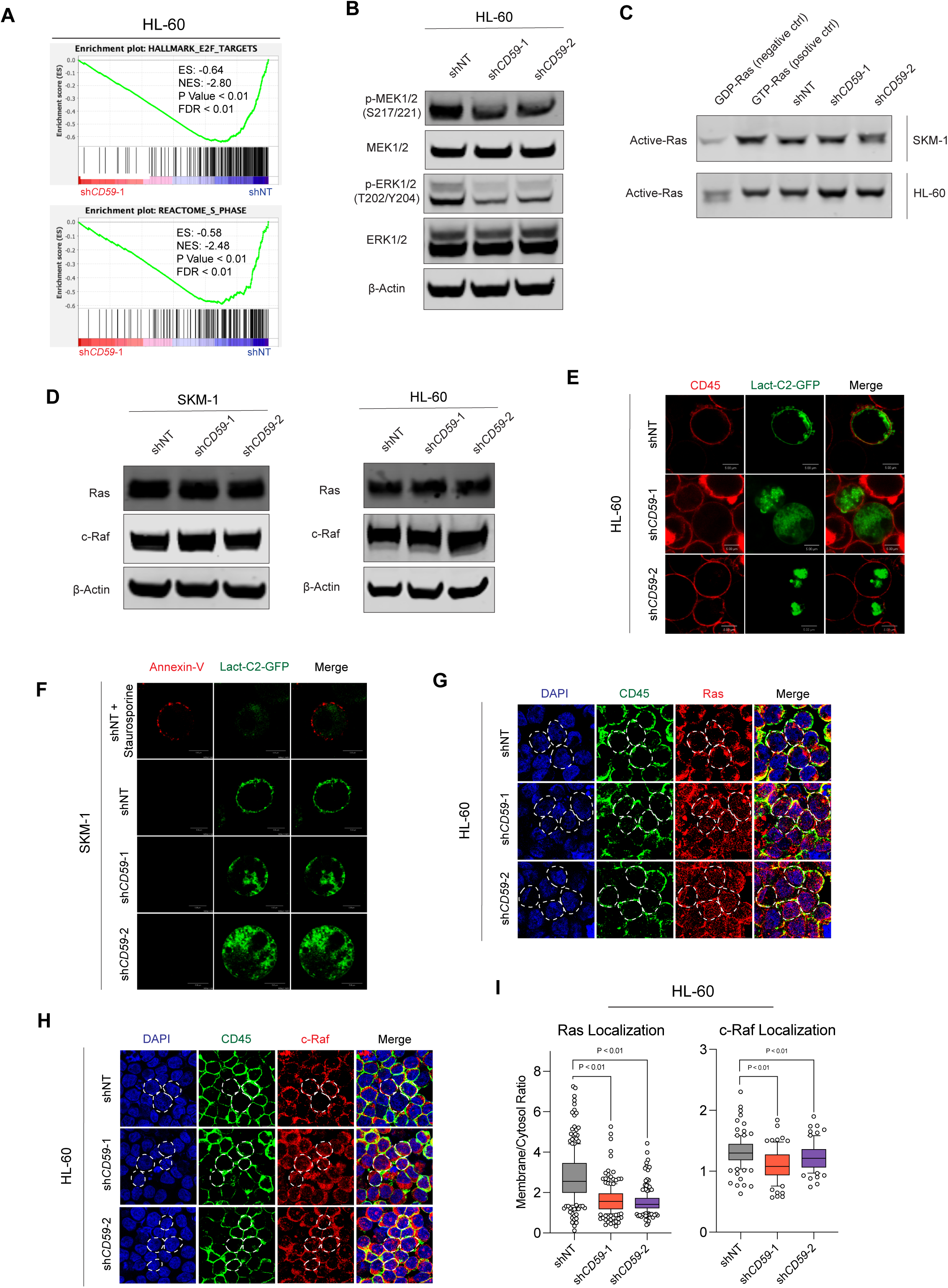
Supporting data for ERK signaling impairment and membrane reorganization following *CD59* silencing in HL-60. **(A)** GSEA enrichment plots for HALLMARK E2F targets and REACTOME S PHASE in sh*CD59*-1 versus shNT-expressing HL-60 cells. **(B)** Western blot for p-MEK (S217/221), MEK1/2, p-ERK1/2 (T202/Y204), ERK1/2, and β-actin in shNT, sh*CD59*-1, and sh*CD59*-2-expressing HL-60 cells, confirming reduced MEK/ERK activation. **(C)** GTP-Ras pulldown assay in SKM-1 (top) and HL-60 (bottom) cells comparing active Ras levels across GDP-Ras (negative control), GTP-Ras (positive control), shNT, sh*CD59*-1, and sh*CD59*-2 conditions, showing no significant change in total Ras activity following *CD59* silencing. **(D)** Western blot for Ras, c-Raf, and β-actin in shNT, sh*CD59*-1, and sh*CD59*-2-expressing SKM-1 (left) and HL-60 (right) cells. **(E)** Representative live cell images of shNT-, sh*CD59*-1, and sh*CD59*-2-expressing HL-60 cells electroporated with Lact-C2-GFP (green) and co-stained with CD45 (red) as an outer membrane marker. White scale bars, 5 μm. **(F)** Representative confocal images of shNT- and sh*CD59*-expressing SKM-1 cells co-stained with Annexin V (red) and Lact-C2-GFP (green). White scale bars, 5 μm. **(G)** Representative confocal images of shNT, sh*CD59*-1, and sh*CD59*-2-expressing HL-60 cells stained with DAPI (blue), CD45 (green), and anti-Ras antibody (red), showing reduced membrane Ras localization in sh*CD59*-expressing cells. White scale bars, 10 μm. **(H)** Representative confocal images of shNT, sh*CD59*-1, and sh*CD59*-2-expressing HL-60 cells stained with DAPI (blue), CD45 (green), and anti-c-Raf antibody (red), showing reduced membrane c-Raf localization in sh*CD59*-expressing cells. White scale bars, 10 μm. **(I)** Quantification of membrane-to-cytosol ratio for Ras and c-Raf in shNT, sh*CD59*-1, and sh*CD59*-2-expressing HL-60 cells (p<0.01).

**Extended Figure 4.**
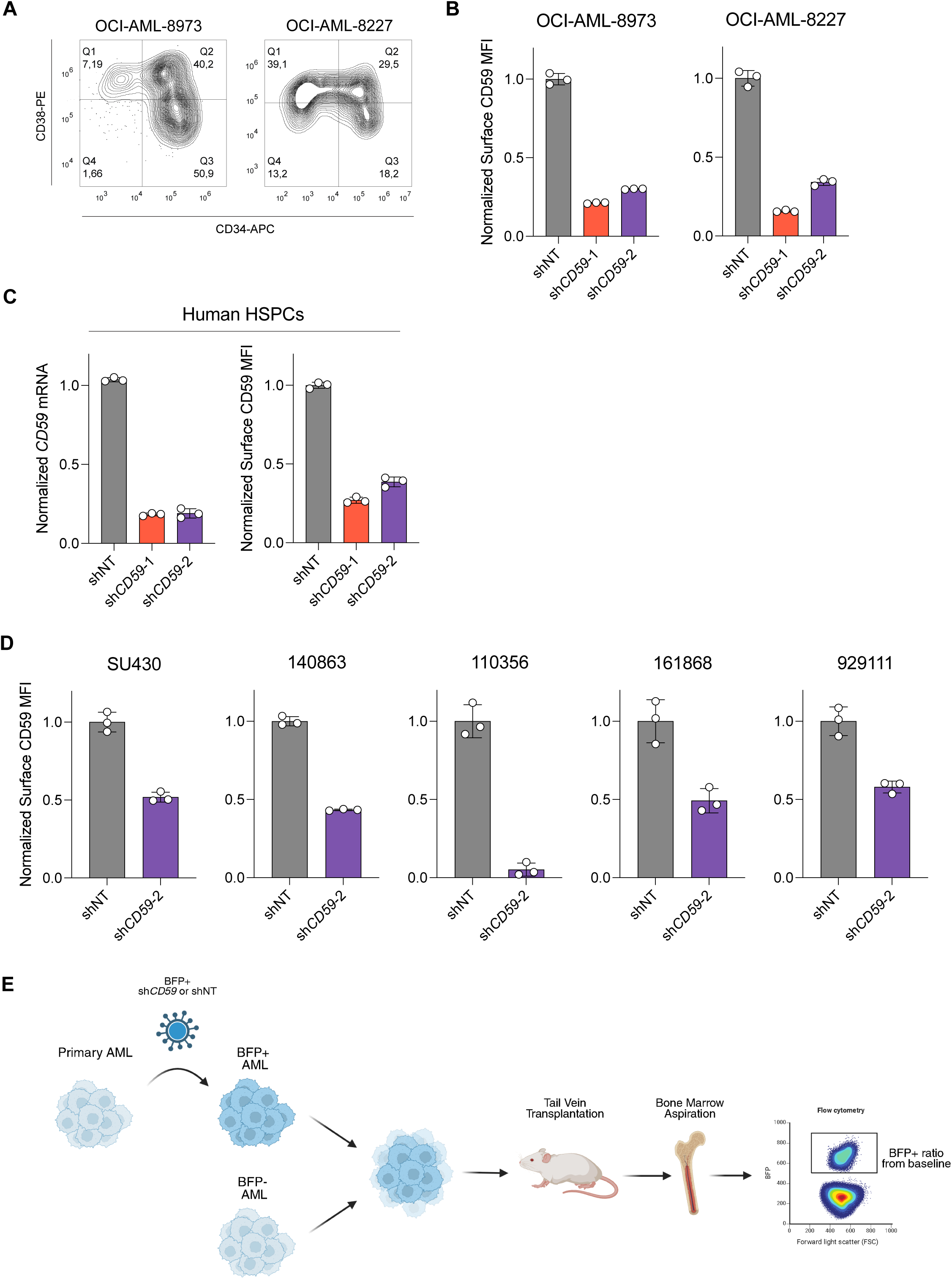
Supporting characterization for LSC and primary AML engraftment experiments. **(A)** Flow cytometry contour plots showing CD34 and CD38 expression in OCI-AML-8973 (left) and OCI-AML-8227 (right), defining the leukemic hierarchy used for CD59 expression analysis. **(B)** Surface CD59 protein expression in OCI-AML-8227 and OCI-AML-8973 using flow cytometry. **(C)** *CD59* mRNA (left) and surface CD59 protein (right) expression in shNT-, sh*CD59*-1-, and sh*CD59*-2-expressing cord blood-derived human HSPCs. **(D)** Surface CD59 protein expression in SU430, 140863, 110356, 161868, 929111 using flow cytometry **(E)** Schematic showing the pipeline on patient-derived xenograft experiments.

**Extended Figure 5.**
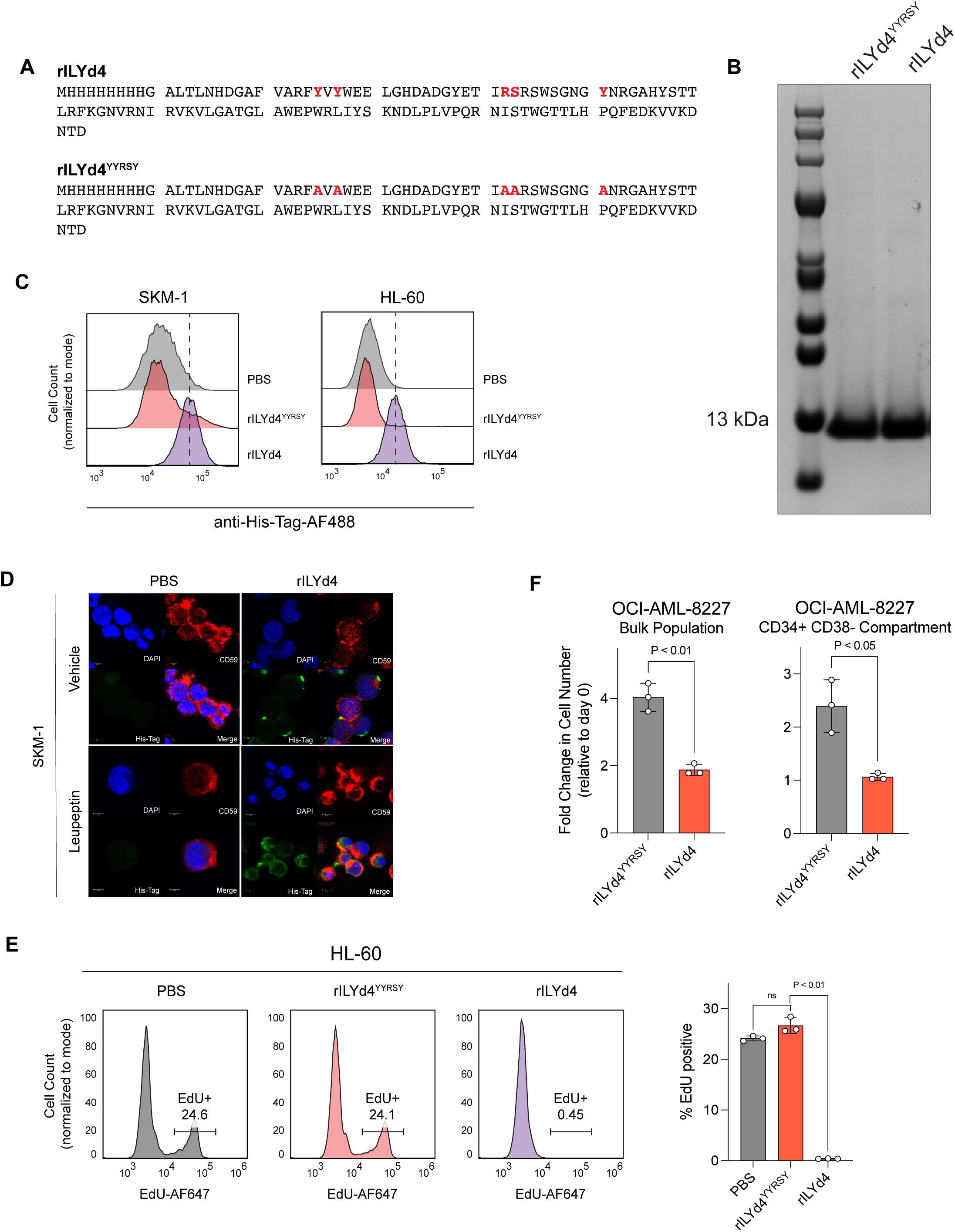
Supporting characterization of rILYd4 production, on-target impact, and CD59-degradation. **(A)** Amino-acid sequences of histidine-tagged rILYd4 and rILYd4^YYRSY^, with the residues important for CD59 binding in rILY4 which were mutated to alanine bolded in red. **(B)** Coomassie-stained SDS-PAGE gel of purified rILYd4 and rILYd4^YYRSY^ showing pure protein at the expected molecular weight. **(C)** Flow cytometry histograms showing binding of histidine-tagged rILYd4^YYRSY^ and rILYd4 to SKM-1 (left) and HL-60 (right) cells detected by anti-His-Tag Alexa Fluor 488. **(D)** Confocal microscopy images of PBS and rILYd4 treated cells treated with vehicle or leupeptin showing lysosomal-mediated CD59 internalization. **(E)** Cell-cycle analysis by EdU incorporation in PBS, rILYd4^YYRSY^, and rILYd4-treated HL-60 cells. **(F)** Fold change in cell number in the bulk population (left) and CD34^+^CD38^-^ LSC-enriched compartment (right) of OCI-AML-8227 following treatment with rILYd4^YYRSY^ or rILYd4 (p<0.01).

**Extended Table 1:** Summary of the mutational and cytogenetic profile of AML cell lines used in this chapter (data curated from Cellosaurus).

| Cell Line | Cytogenetics | Mutations |
| --- | --- | --- |
| SKM-1 | Hypodiploid karyotype (CK) | *TP53* R248Q, *ASXL1*, *KRAS* p.K117N |
| HL-60 | Deletion of chr. 17, chr. 8, 1 arm of chr. 16 (CK) | *TP53* del., *NRAS* p.Q61L |
| NB-4 | Hypertriploid karyotype (CK) | *TP53* R248Q |
| EOL-1 | Hyperdiploid karyotype (CK) | *MT2A* (*MLL*) partial tandem duplication |
| THP-1 | Hyperdiploid karyotype (CK) | *KMT2A-MLLT3* fusion<br>*TP53* R174fs*3 |
| KG-1α | Hyperdiploid karyotype (CK) | *NRAS* G12D<br>*TP53* c672+1G>A |
| MOLM-13 | Hyperdiploid karyotype | *KMT2A-MLLT3* fusion<br>*FLT3*-ITD |
| OCI-AML-3 | Hyperdiploid karyotype | *DNMT3A* R882C, *NRAS* G61L, *NPM1* W288C |
| NOMO-1 | Hypertriploid karyotype (CK) | *KMT2A-MLLT3*, *KRAS* G13D,<br>*TP53* del. |

**Extended Table 2:** Summary of the mutational and cytogenetic profile of primary AML cells.

| Sample ID | ELN2022 | Cytogenetics | NGS |
| --- | --- | --- | --- |
| 161868 | Intermediate | Normal karyotype | *ETV6*, *FLT3*, *NPM1*,<br>*DNMT3A* |
| 929111 | Adverse | 46~51,XY ,add(5)(q11.2),-7,-9,-11,inv(12)(p13q24.1),?del(13(q22),-17,-21,+r,+3~6mar[cp11] | *TP53* |
| 140863 | Adverse | 42~45,XY,-3,-5,del(7)(q21),der(12)t(12;?16)(p13;?11.2),-13,-14,-16,-20,+r,+4mar[cp10] | *ASXL1*, *CALR*,<br>*CSF3R*, *IDH1*,<br>*SETBP1*, *SRSF2* |
| 110356 | Adverse | 44,XX,inv(3)(q21q26),del(5)(q13q33),-7,add(8)(p21),t(9;13)(q32 or q34;q14),?dic(12;22)(p13;q13)[9]/44,idem,?del(4)(q31)[4] | NSG not performed |
| SU430 | Adverse | Der(7)t(7;11)(q11.2;q13) | *IDH1*-R132C,<br>*DNMT3A*-R882C |

**Extended Table 3:**
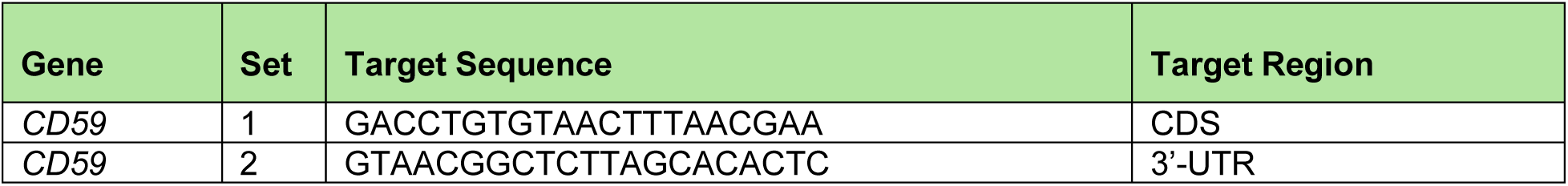
shRNA sequences used for *CD59* silencing.

**Extended Table 4:**
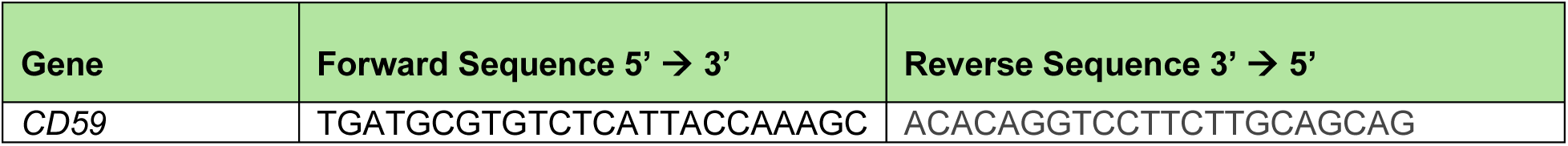
RT-qPCR primers used to measure *CD59* mRNA levels.

**Extended Table 5:** List of antibodies and dyes used for flow cytometry.

| Target | Host Species | Dilution | Manufacturer | Catalog No. |
| --- | --- | --- | --- | --- |
| Human CD59 | Mouse | 1:100 | Abcam | ab9183 |
| Human CD34 | Mouse | 1:100 | Biolegend | 343509 |
| Human CD38 | Mouse | 1:100 | Biolegend | 356603 |
| Human CD45 | Mouse | 1:20 | Biolegend | 304011 |
| Mouse CD45.1 | Mouse | 1:100 | Biolegend | 110751 |
| Mouse TER-199 | Rat | 1:100 | Biolegend | 116205 |
| Mouse IgG2b | Mouse | 1:100 | BioRad | MCA691 |
| Anti-mouse IgG (H+L) | Goat | 1:750 | Cell Signaling Technology | 4410 |
| APC annexin V | N/A | 1:200 | Biolegend | 640941 |
| SYTOX green | N/A | 1:1000 | Thermo fisher | S34860 |
| His-tag | Rabbit | 1:100 | Cell Signaling Technology | 2365 |

**Extended Table 6:** List of antibodies used for Western Blots.

| Target | Host Species | Dilution | Manufacturer | Catalog No. |
| --- | --- | --- | --- | --- |
| Human CD59 | Rabbit | 1:10,000 | Cell Signaling Technology | 25393 |
| Human ERK1/2 | Rabbit | 1:10,000 | Cell Signaling Technology | 4695 |
| Human p-ERK1/2 (T202,Y204) | Rabbit | 1:10,000 | Cell Signaling Technology | 4370 |
| Human MEK1/2 | Rabbit | 1:10,000 | Cell Signaling Technology | 9122 |
| Human p-MEK1/2 (S217/221) | Rabbit | 1:10,000 | Cell Signaling Technology | 9154 |
| Human beta-Actin | Mouse | 1:10,000 | Cell Signaling Technology | 3700 |
| Human Ras | Mouse | 1:200 | Cell Signaling Technology | 8821 |

**Extended Table 7:** List of antibodies and dyes used for confocal imaging.

| Target | Host Species | Dilution | Manufacturer | Catalog No. |
| --- | --- | --- | --- | --- |
| Human CD59 | Mouse | 1:100 | Abcam | ab9183 |
| Human CD45 | Mouse | 1:100 | Proteintech | 4E9B2 |
| Human Ras | Rabbit | 1:100 | Abcam | ab108602 |
| Human c-Raf | Rabbit | 1:100 | Invitrogen | MA5-53558 |
| His-tag | Rabbit | 1:100 | Cell Signaling Technology | 2365 |
| Anti-mouse IgG (H+L) | Goat | 1:750 | Cell Signaling Technology | 4410 |
| Anti-rabbit IgG (H+L) | Goat | 1:750 | Cell Signaling Technology | 4412 |
| DNA (Antifade + DAPI) | N/A | N/A | Cell Signaling Technology | 8961 |

**Extended Table 8:**
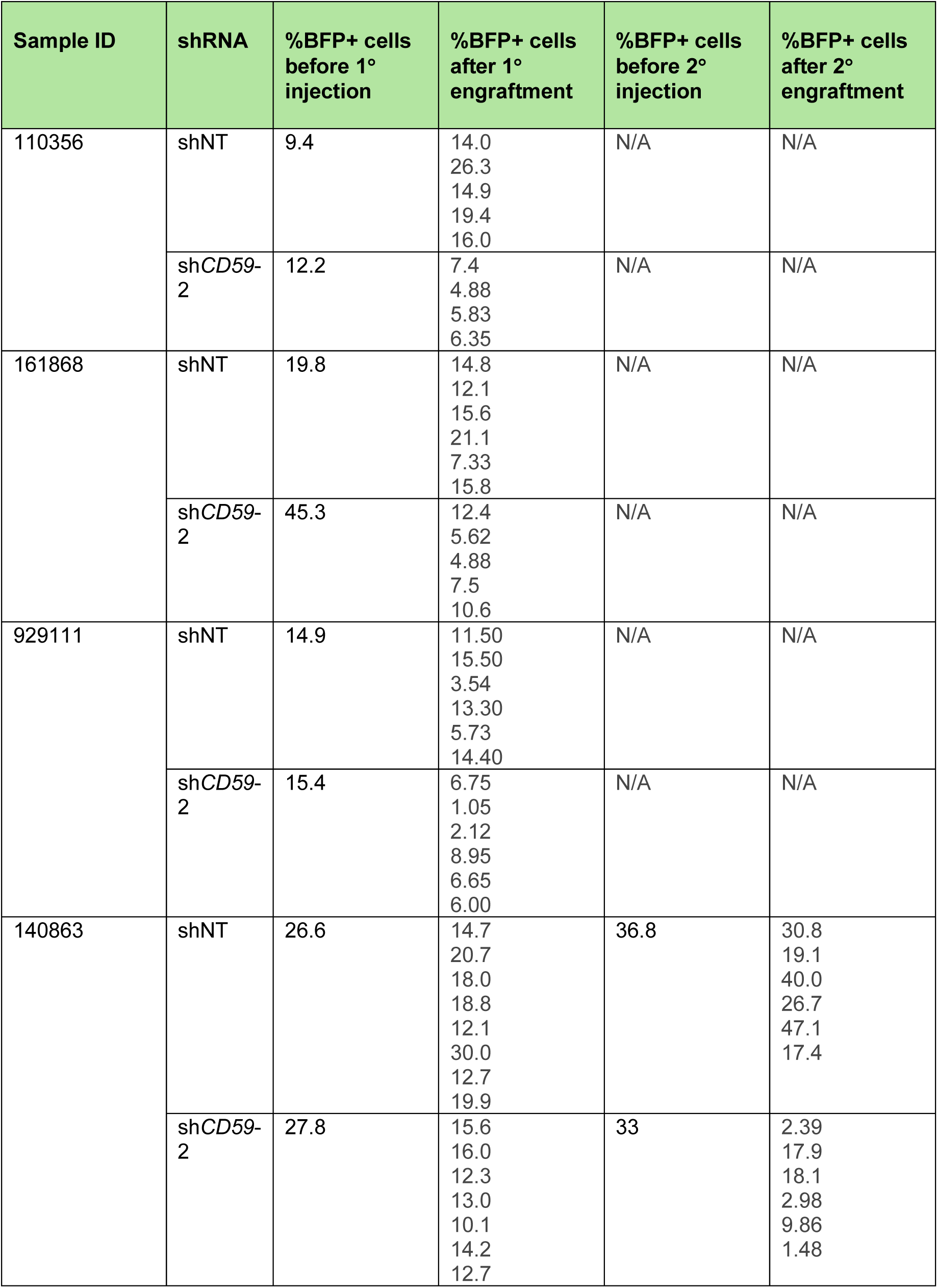
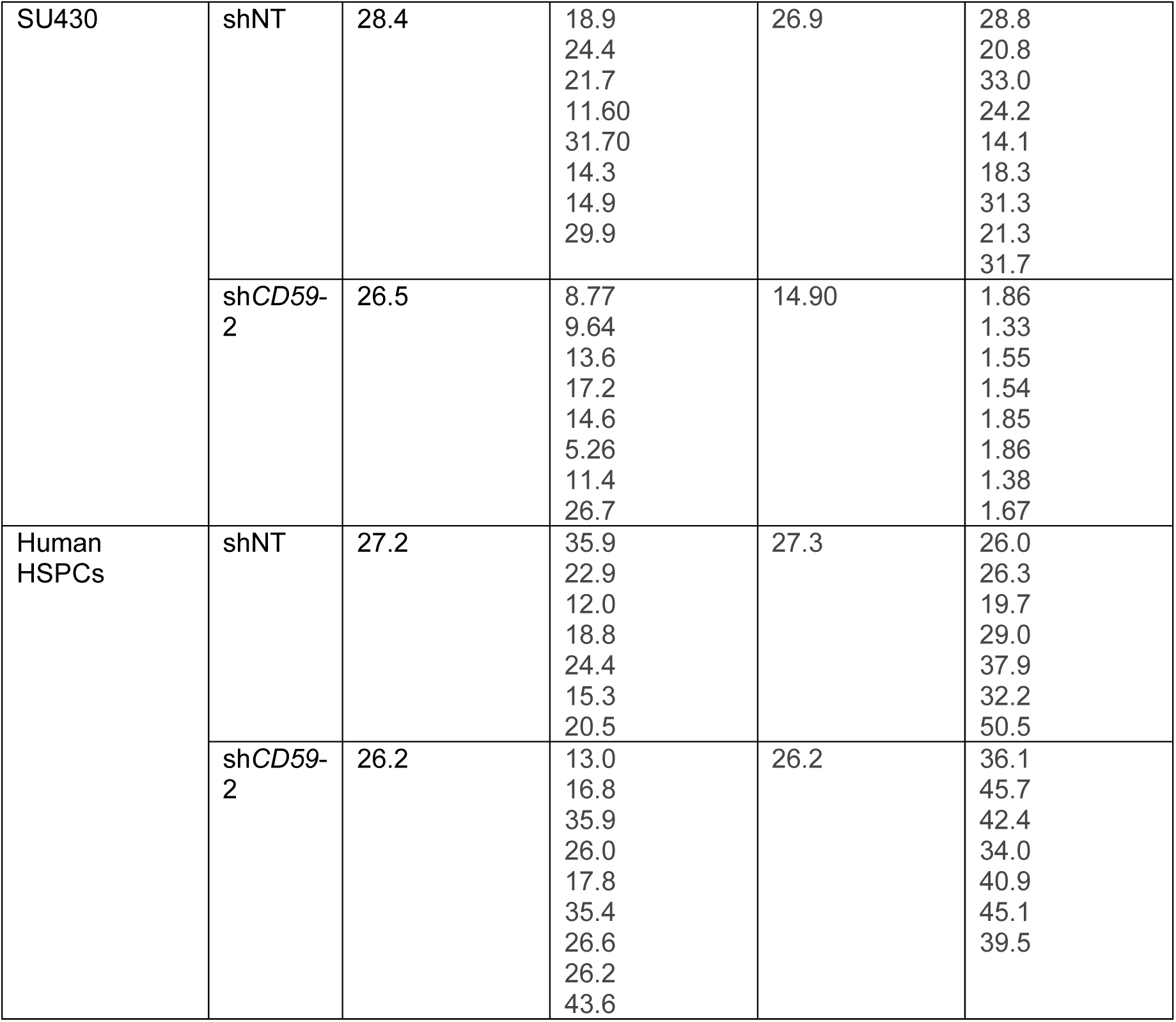
Percentage of BFP+ primary AML samples and human HSPCs at baseline and at the time of experimental endpoint.

| Sample ID | shRNA | %BFP+ cells before 1° injection | %BFP+ cells after 1° engraftment | %BFP+ cells before 2° injection | %BFP+ cells after 2° engraftment |
| --- | --- | --- | --- | --- | --- |
| 110356 | shNT | 9.4 | 14.0<br>26.3<br>14.9<br>19.4<br>16.0 | N/A | N/A |
|  | shCD59-2 | 12.2 | 7.4<br>4.88<br>5.83<br>6.35 | N/A | N/A |
| 161868 | shNT | 19.8 | 14.8<br>12.1<br>15.6<br>21.1<br>7.33<br>15.8 | N/A | N/A |
|  | shCD59-2 | 45.3 | 12.4<br>5.62<br>4.88<br>7.5<br>10.6 | N/A | N/A |
| 929111 | shNT | 14.9 | 11.50<br>15.50<br>3.54<br>13.30<br>5.73<br>14.40 | N/A | N/A |
|  | shCD59-2 | 15.4 | 6.75<br>1.05<br>2.12<br>8.95<br>6.65<br>6.00 | N/A | N/A |
| 140863 | shNT | 26.6 | 14.7<br>20.7<br>18.0<br>18.8<br>12.1<br>30.0<br>12.7<br>19.9 | 36.8 | 30.8<br>19.1<br>40.0<br>26.7<br>47.1<br>17.4 |
|  | shCD59-2 | 27.8 | 15.6<br>16.0<br>12.3<br>13.0<br>10.1<br>14.2<br>12.7 | 33 | 2.39<br>17.9<br>18.1<br>2.98<br>9.86<br>1.48 |
| SU430 | shNT | 28.4 | 18.9<br>24.4<br>21.7<br>11.60<br>31.70<br>14.3<br>14.9<br>29.9 | 26.9 | 28.8<br>20.8<br>33.0<br>24.2<br>14.1<br>18.3<br>31.3<br>21.3<br>31.7 |
|  | shCD59-2 | 26.5 | 8.77<br>9.64<br>13.6<br>17.2<br>14.6<br>5.26<br>11.4<br>26.7 | 14.90 | 1.86<br>1.33<br>1.55<br>1.54<br>1.85<br>1.86<br>1.38<br>1.67 |
| Human HSPCs | shNT | 27.2 | 35.9<br>22.9<br>12.0<br>18.8<br>24.4<br>15.3<br>20.5 | 27.3 | 26.0<br>26.3<br>19.7<br>29.0<br>37.9<br>32.2<br>50.5 |
|  | shCD59-2 | 26.2 | 13.0<br>16.8<br>35.9<br>26.0<br>17.8<br>35.4<br>26.6<br>26.2<br>43.6 | 26.2 | 36.1<br>45.7<br>42.4<br>34.0<br>40.9<br>45.1<br>39.5 |

